# DNA Transposons favour de *novo* transcript emergence through enrichment of transcription factor binding motifs

**DOI:** 10.1101/2023.10.03.560692

**Authors:** Marie Kristin Lebherz, Bertrand Fouks, Julian Schmidt, Erich Bornberg-Bauer, Anna Grandchamp

## Abstract

*De novo* genes emerge from non-coding regions of genomes via succession of mutations. Among others, such mutations activate transcription and create a new open reading frame (ORF). Although the mechanisms underlying ORFs emergence are well documented, relatively little is known about the mechanisms enabling new transcription events. Yet, in many species a continuum between absent and very prominent transcription has been reported for essentially all regions of the genome.

In this study we searched for *de novo* transcripts by using newly assembled genomes and transcriptomes of seven inbred lines of *Drosophila melanogaster*, originating from six European and one African population. This setup allowed us to detect line specific *de novo* transcripts, and compare them to their homologous non-transcribed regions in other lines, as well as genic and intergenic control sequences. We studied the association with transposable elements and the enrichment of transcription factor motifs upstream of *de novo* emerged transcripts and compared them with regulatory elements.

We found that *de novo* transcripts overlap with TEs more often than expected by chance. The emergence of new transcripts correlates with high CpG islands and regions of TEs activity. Moreover, upstream regions of *de novo* transcripts are highly enriched with regulatory motifs. Such motifs abound in new transcripts overlapping with TEs, particularly DNA TEs, and are more conserved upstream *de novo* transcripts than upstream their non-transcribed homologs. Overall, our study demonstrates that TEs insertion is important for transcript emergence, partly by introducing new regulatory motifs from DNA TE families.

## Introduction

For long, new genes were thought to exclusively arise from pre-existing genes (Guerzoni and McLysaght, 2011). However recent studies showed that a non-negligible proportion of new genes also emerge *de novo* from non-coding regions of the genome (Schlötterer, 2015; Bornberg-Bauer et al., 2015; Rödelsperger et al., 2019; Tautz and Domazet-Lošo, 2011; McLysaght and Hurst, 2016; Van Oss and Carvunis, 2019; Bornberg-Bauer et al., 2021). Several *de novo* genes have been shown to become essential, bearing important organismal functions,e.g. male fertility (Gubala et al., 2017) and cold resistance (Baalsrud et al., 2018). For a *de novo* gene to arise, it requires both the gain of an open reading frame (ORF) and the acquisition of transcription (Durand et al., 2019; Schlötterer, 2015). While the gain of ORFs in the emergence of *de novo* genes has been well studied (Zhuang and Cheng, 2021; Delihas, 2022; Rödelsperger et al., 2019; Wang et al., 2020b; Carvunis et al., 2012; Grandchamp et al., 2023b), how transcription is acquired remains poorly understood.

The transcription of a gene is initiated at the core promoter which is located upstream the gene’s 5’ untranslated region (UTR) (Haberle and Stark, 2018; Butler and Kadonaga, 2002). Core promoters contain specific binding motifs, such as the *TATA box* or the *Initiator sequence*, that are recognized by transcription factors (tFs) (Boeva, 2016). Binding motifs with low identity to the consensus sequence are referred as minimal motif (Wang et al., 2020a). Transcription factors then recruit the protein complexes required for transcription (Butler and Kadonaga, 2002). However, transcription of low amounts of transcripts can also be initiated by a core promoter alone (reviewed in Haberle and Stark (2018); Small and Arnosti (2020)). Promoters can also produce antisense transcripts by initiating transcription in both direction (Scruggs et al., 2015). Furthermore, proximal and distal enhancers regulate the levels of transcription. Proximal enhancers (also called proximal promoters) are located directly upstream of core promoters, while distal enhancers influence transcription over long distances (Kim and Shiekhattar, 2015; Haberle and Stark, 2018). Both contain tF binding motifs and can increase the amount of transcription initiated by the promoter (Haberle and Stark, 2018), independently of their locations and directions (Haberle and Stark, 2018). Enhancers often carry out bi-directional transcription, producing short but unstable transcripts in both directions (Small and Arnosti, 2020; Meers et al., 2018). Enhancers and promoters can also occasionally be converted into each other (Majic and Payne, 2020), and promoters can be interconnected by successive mutations without completely loosing their activity (Kurafeiski et al., 2019)

In a non-coding region, the gain of transcription can result from random point mutations in a minimal motif and lead to stable transcription (Palazzo and Lee, 2015; Kapusta and Feschotte, 2014), as genomes generally contain many cryptic functional sites with minimal promoters (Kapusta and Feschotte, 2014). Genomic mutations can also be initiated via the insertion of transposable elements (TEs). TEs are mobile DNA sequences that can move and amplify in genomes. They can be divided into two classes, based on their transposition mode: RNA and DNA transposons, which are further divided into sub classes and families based on their sequence characteristics (McCullers and Steiniger, 2017). Several studies reported major reshuffling of genomic architectures due to TEs, as well as their role in adaptive evolution (Bourque et al., 2018; Delprat et al., 2009; Thybert et al., 2018). For example, syncytin genes, enabling cell-cell fusion in mammalian placenta, are derived from TEs (Malik, 2012). TEs have also aided the evolution of the placenta in mammals, by acting on enhancers activity (Chuong et al., 2013). Other epigenetic mechanisms can influence transcription levels, such as DNA methylation, which represses genes transcription in vertebrates via the modulation of tFs activity (Law and Jacobsen, 2010). In invertebrates, methylation patterns are also associated with the regulation of transcription (Dixon and Matz, 2021), but the correlation between transcription and methylation is less clear than in vertebrates (Dunwell and Pfeifer, 2014; Lyko et al., 2000). Transcription is a highly dynamic and plastic process with high rates of transcripts gain and loss in closely related species, as well as among populations and individuals (Zhao et al., 2014; Grandchamp et al., 2022, 2023a; Neme and Tautz, 2016; Iyengar and Bornberg-Bauer, 2023), suggesting fast transcripts turnover. However, the mechanisms promoting *de novo* transcripts, i.e. transcription initiation from non-coding regions, remains elusive.

In this study, we investigate the mechanisms underlying novel transcript emergence at short evolutionary time scales by studying *de novo* transcripts in seven lines of *Drosophila melanogaster*, originating from different geographical locations (Grandchamp et al., 2022). By using long-read sequencing and a common annotation methodology across all genomes, our genomes and transcriptomes present a unique opportunity to precisely categorize *de novo* transcripts in each *Drosophila* line and investigate the molecular basis underlying the gain of transcription. Indeed, our dataset is allowing us to compare directly the related DNA sequences that are transcribed in one or several lines but not in others. In particular, we studied the role of transposable element insertions and motif enrichment upstream of *de novo* transcripts that emerged in each *Drosophila* line. Overall, our analyses reveal that the emergence of transcription is aided by an enrichment of motifs upstream of a DNA sequence, motif enrichment which is itself favored by nearby insertion of DNA transposons.

## Results

### General characteristics of de novo transcripts

To characterize the molecular basis underlying gains of transcription, we used a conservative approach to define *de novo* transcripts, ensuring detection of strictly *de novo* transcript (see Methods). Such definition and filtering led to the discovery of between 403 (Sweden [SE]) to 628 (Ukraine [UA]) *de novo* transcripts across *D. melanogaster* lines (mean = 504 *±* 28.04 (SE), Figure 1 A, Supplemental deposit). *De novo* transcripts were unevenly distributed among and along chromosomes, with the highest numbers of *de novo* transcripts in 3L and 3R chromosome arms (SI-S1). Most of *de novo* transcripts were found in only one *D. melanogaster* line (2389 / 3528) and only a few (38) were shared among all lines, suggesting a high birth / death rate of *de novo* transcripts, Figure 1 B). This high birth/death rate of de novo transcripts is likely the result of gain / loss of transcription, as most *de novo* transcripts (14058 blast hit out of 18903 blast searches in a maximum of 6 lines) had a ‘non-transcribed’ homolog in at least one other *D. melanogaster* line (Supplemental deposit). Moreover, *de novo* transcripts show different patterns from annotated transcripts (both genes and non-coding RNAs), with *de novo* transcripts having lower expression level, GC content, exon number, and spliced length compared to annotated transcripts (GLMM: TPM: p <0.001, GC content: p <0.001, exon number: p <0.001, spliced length: p <0.001, (SI-S2)).

**Figure 1:**
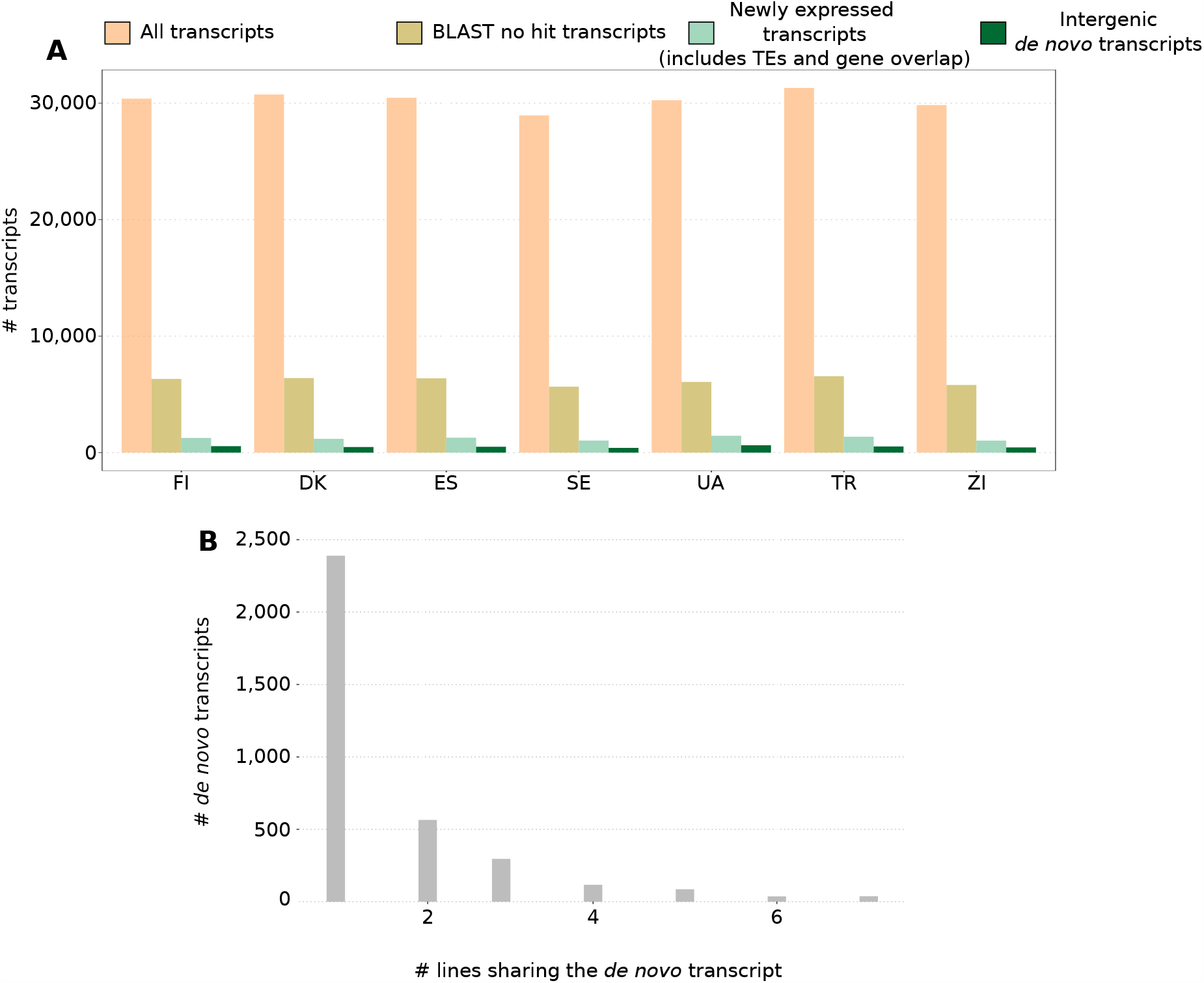
*De novo* transcripts. (A): Number of transcripts after filtering steps. The beige bar represents all transcripts detected with transcriptome assembly. The grey bar represents all transcripts without a BLAST hit. The green bar represents *de novo* transcripts after filtering for TPM and splicing. The dark green bar represents only the intergenic *de novo* transcripts after filtering out transcribed TE. (B): Number of *de novo* transcripts shared by lines

### DNA transposon insertions favour the gain of transcription

For each genome, we performed a *de novo* annotation of TEs, using the TransposonUltimate pipeline (Riehl et al., 2022) (method, Supplemental deposit). To understand how TEs can favour the gain of transcription, we first assessed the relationship between TEs and *de novo* transcripts at the chromosome scale (Figure 2). While *de novo* transcripts were evenly distributed along chromosomes, inactive TEs and expressed TEs, were in higher density in the telomere regions of chromosomes (GLMM: p <0.001; Figure 2 A, SI-S3). Nevertheless, *de novo* transcript densities were positively correlated with TE densities at a 100 kb scale (GLMM: p <0.001). An important mechanism by which TE impact gene expression is the import of epigenetic marks, such as DNA methylation (Zhou et al., 2020). We therefore calculated the CpGoe, as an estimate for DNA methylation status, with high CpGoe values corresponding to low level of methylation. *De novo* transcripts displayed low level of methylation (CpGoe: mean & median = 0.902 +-sd 0.222) and their methylation stati were negatively correlated with TE density (GLMM: p <0.001), highlighting the role of TEs in importing epigenetic marks (Figure 2 B, Supplemental deposit, SI-S4).

**Figure 2:**
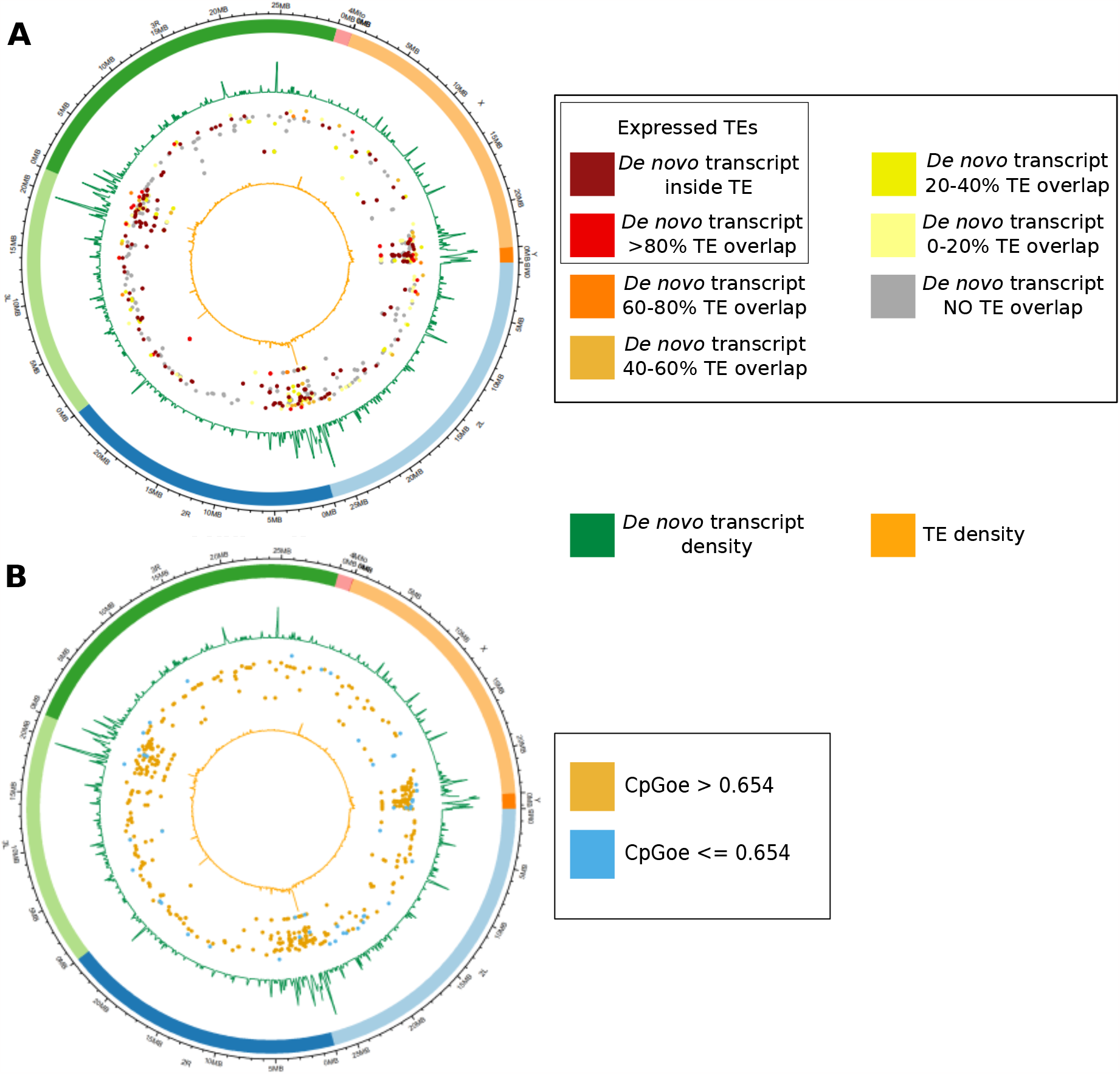
*De novo* transcripts and TE density among the chromosomes. The circular plots represent the *D*.*melanogaster* line collected in Denmark (DK). Plots with similar distributions can be found for all other lines in the *supplemental data*. The 8 chromosome arms are represented with specific colours. In the 2 circle plots, the green circles represent *de novo* transcripts, and the yellow lines represent TEs distribution. (A) The coloured dot represents expressed TEs and *de novo* transcripts distribution according to their relative overlap with TEs. (B) The coloured dots represent the CpGoe values of *de novo* transcripts according to their genomic distribution.

In addition to our chromosome scale analyses, we also investigated the impact of TE insertions on *de novo* transcripts by comparing the number of TE overlapping with these transcripts, as well as their down- and upstream regions, with random intergenic regions as a negative control. *De novo* transcripts displayed a higher amount of TE insertions compared to other sequences, however with a lower length of TE overlap (GLMM: p <0.001; Figure 3 A,B, Supplemental deposit). Furthermore to be able to pinpoint precisely the role of TE insertions on the gain of transcription, we directly compared *de novo* transcripts with their ‘non-transcribed’ homolog sequences present in other *D. melanogaster* lines. Our analyses revealed that TE insertions did not differ between *de novo* transcripts and ‘non-transcribed’ homologs, however *de novo* transcripts displayed shorter overlaps with TEs as well as a lower number of TE insertions compared to ‘non-transcribed’ homologs (GLMM, p <0.001, SI-S5). Moreover, RNA TEs were less abundant in *de novo* transcripts compared to ‘non-transcribed’ homologs (GLMM, p < 0.001,Figure 3 C, SI-S5). On the contrary, DNA TEs were more abundant in *de novo* transcripts compared to ‘non-transcribed’ homologs (GLMM, p < 0.001). Our results highlight a different impact between TE classes (DNA vs. RNA) on transcription gain.

**Figure 3:**
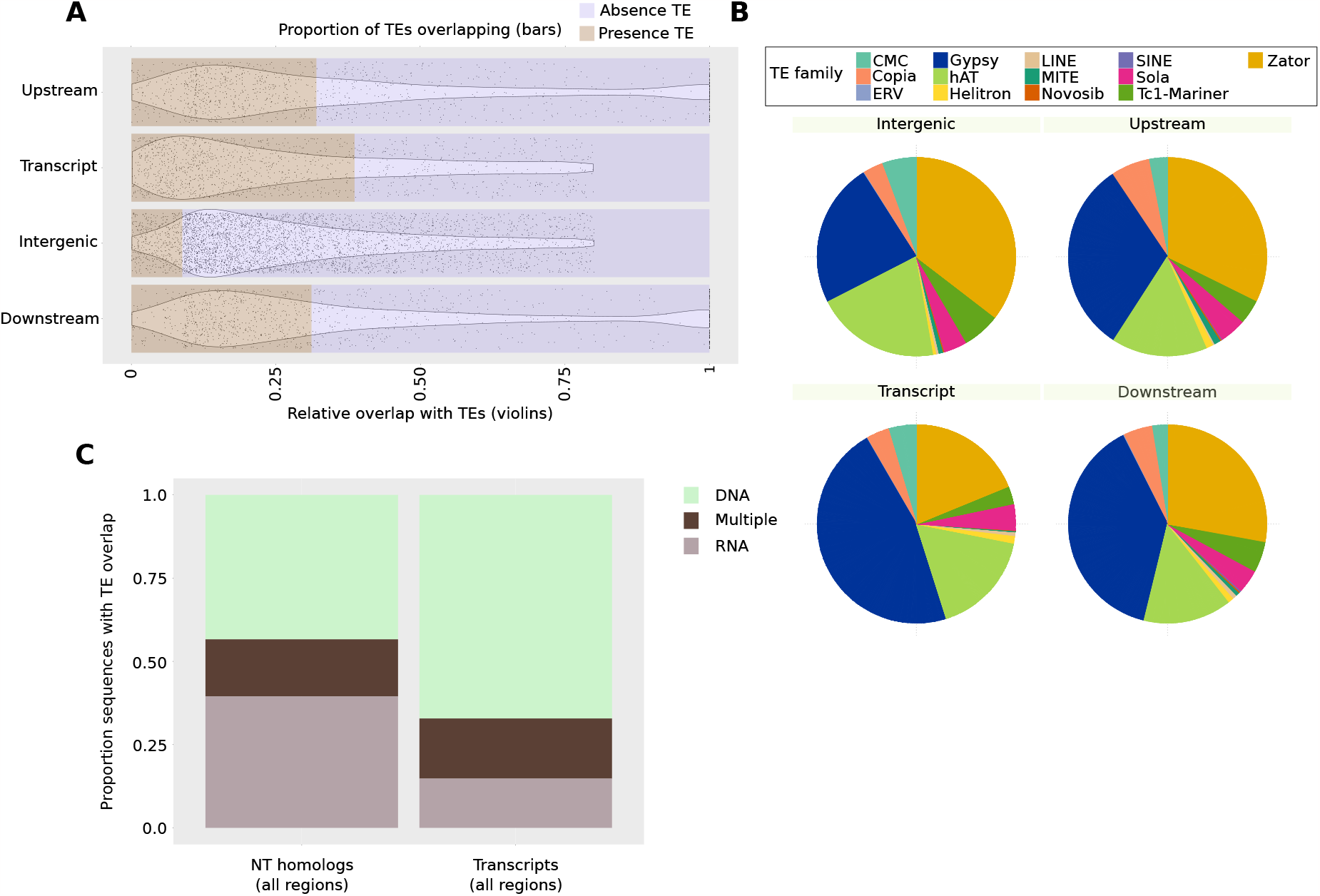
TEs overlap. (A) Relative sequence overlap with TEs and number of sequences overlapping with TEs into four datasets: Intergenic sequences, upstream sequences of *de novo* transcripts, downstream sequences of *de novo* transcripts, *de novo* transcripts. (B) Percentage of TEs overlapping with the four datasets according to their families.(C) Major classes of TEs overlapping with *de novo* transcripts and their non-transcribed homologs.

### Motifs enrichment

A major factor influencing gene expression is the presence of specific DNA motifs enabling the transcription machinery to bind to the DNA region. We therefore investigated the role of DNA binding motifs for the gain of transcription. We compared several measures of motif enrichment (specific to both tF motifs from enhancers and distal promoters, as well as (core) promoters) upstream of our *de novo* transcripts, as positive controls upstream of genes and expressed TEs, and as negative control random intergenic regions. Motif enrichments were further divided into two classes according to their thresholds of similarity to their PSSM matrix : low identity motifs (minimal motifs), with a score of identity to the matrix > 80%, and high identity motifs, with an ID score of 95% identity as a minimum (Figure 4). This comparison revealed that TEs and *de novo* transcripts overlapping with TEs have higher number of low identity tF motifs compared to other sequences (GLMM: p <0.001). Moreover, *de novo* transcripts that do not overlap with TEs displayed higher numbers of core promoters with high identity score than TEs and *de novo* transcripts (GLMM: p <0.001; SI-S6,S7). Overall, genes and intergenic regions displayed a higher enrichment of core promoter motifs (both high and low identity motifs) and of tF motifs with high identity score, while TEs and *de novo* transcripts displayed an enrichment of tF motifs of low identity score (GLMM, p <0.001,Figure 4, SI-S6,S7).

**Figure 4:**
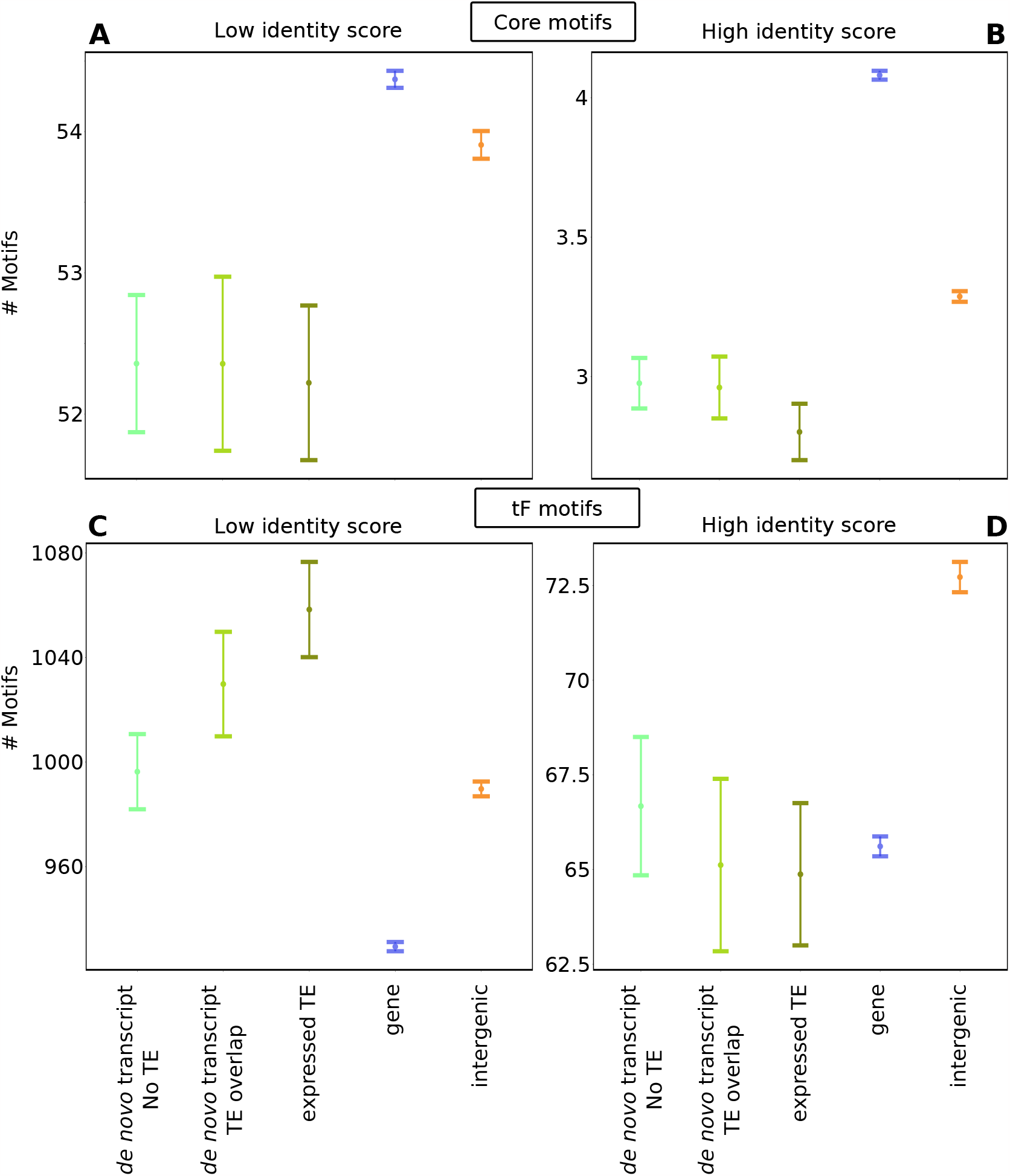
Number of motifs detected upstream five sequences datasets. (A) Number of low identity Core promotor (0.8) motifs detected upstream i) *de novo* transcripts overlapping no TE (light green), ii) *de novo* transcripts overlapping with TEs (green), iii) expressed TEs (dark green) iv) genes (blue), v) randomly selected intergenic regions that are not transcribed (orange). (B) Number of high identity Core promotor motifs (0.95) detected upstream the aforementioned dataset of sequences (C) Number of low identity tF motifs (0.80) detected upstream the aforementioned dataset of sequences. (D) Number of high identity tF motifs (0.95) detected upstream the aforementioned dataset of sequences.

When studying motifs individually (SI-S6,S7), 13 motifs were enriched upstream *de novo* transcripts and TEs, compared to intergenic regions, with a high threshold (relative score = 0.95; supp data), four of them being also significantly enriched in upstream genes. Three of these 13 motifs were significantly enriched upstream *de novo* transcripts without TE overlap. 11 out of these 13 motifs were specific for homeo domain factors, with one zinc finger factors (Supplemental deposit). Among the ten most abundant motifs (ara, mirr, CG4328-RA, lbe, PHDP, H2.0, Deaf.1, caup, C15, lbl), four were enriched in *de novo* transcribed TEs and in TEs overlapping *de novo* transcripts. We found 78 tF motifs that were enriched upstream *de novo* transcripts and TEs with a low threshold (relative score = 0.8), 13 of them being also significantly enriched in genes. Only 18 of them were enriched upstream *de novo* transcripts that did not overlap any TE. Most of these 73 motifs were specific tF for homeo domain factors or zinc finger, however they also included one motifs for high mobility group domain factor, for one heat shock factor, two motifs for leucine zipper factors, two for paired box factors, one fork head/winged helix factor, for a STAT and TEA domain factor. Out of the ten most frequent motifs from the dataset using this treshold (CG4328-RA, br, H2.0, PHDP, C15, vvl, Dbx, ct, lbl, ara) seven were enriched in all *de novo* transcipts, one of them also in genes. Additional two were enriched only in TEs and *de novo* transcripts overlapping TEs.

In addition, we compared directly binding motif enrichment upstream sequences of *de novo* transcripts and their ‘non-transcribed’ homologs. We observed no significant difference in motif enrichment between *de novo* transcripts and their ‘non-transcribed’ homologs. The best statistical model included the enrichment of low identity core promoters but it was not significant (GLMM, p = 0.136, SI-S8). Furthermore, we implemented the impact of TE insertions along motif enrichment between *de novo* transcripts and their ‘non-transcribed’ homologs. *De novo* transcripts exhibit, when TE inserted, a higher density of tF motifs of low identity, suggesting that TE insertions enable transcription through low tF motif enrichment (Figure 5). Finally, we accounted for the different TE class (DNA vs. RNA) inserting among *de novo* transcripts and their non-transcribed homologs. While *de novo* transcripts have a lower ratio of RNA transposons compared to their ‘non-transcribed’ homologs, high number of RNA transposon insertions in *de novo* transcripts is linked with an enrichment low identity core promoter motifs (GLMM, p <0.001, SI-S8).

**Figure 5:**
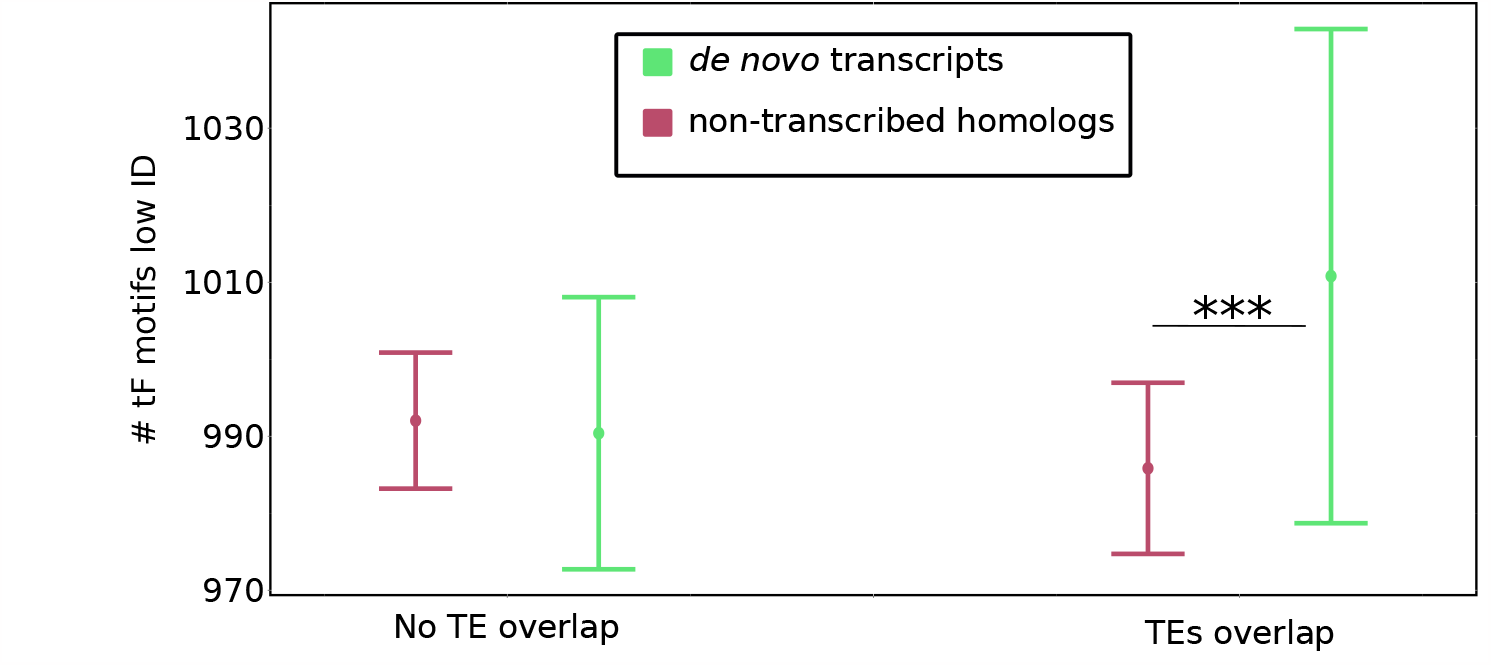
Enrichment in low tF promoter motifs upstream *de novo* transcripts and their non-transcribed homologs. The green colour represents *de novo* transcripts. The pink colour represents non-transcribed homologs. The bars on the left represent sequences without TE overlap, while the bars on the right represent sequences with TE overlap. The y axis represents the number of low tF motifs.

## Discussion

### Detection of de novo transcripts

To understand how transcription can be gained in intergenic regions leading to the emergence of *de novo* genes, we searched for *de novo* intergenic transcripts that emerged in seven lines of *Drosophila melanogaster*. Our stringent definition led to the discovery of 3,799 transcripts over 7 *D. melanogaster* lines, with an average of 504 intergenic *de novo* transcripts per line. This amount of *de novo* transcripts, while being lower than in a previous study of new transcripts emergence in lines (Everett et al., 2020), corresponds well to previous estimates (Camilleri-Robles et al., 2022; Huang et al., 2015), if we account only for intergenic *de novo* transcripts.

Moreover, the characteristics of our *de novo* transcripts corresponds well to those of previous studies, namely a lower expression, lower GC content, lower number of exons, and a shorter sequence than known genes. Finally, our estimation of *de novo* transcripts could have been minimized by not accounting for transcripts with low level of expression or tissue- and life-stage specific expression, resulting in lower detection of *de novo* transcripts (Grandchamp et al., 2022).

### Overlap with transposable elements

Among all detected *de novo* transcripts, 34% overlapped fully or by more than 80% with TEs, and were then considered as “active TEs” rather than *de novo* transcripts. This first outcome suggests that TEs have important mobility inside the species. TEs were massively detected and active in the telomeric regions of the chromosomes, as previously reported (Kordyukova et al., 2018). While *de novo* transcripts display a higher proportion of TE insertions compared to control sequences, TEs overlapped mainly with small fractions of the transcripts and of their surrounding regions, rejecting the hypothesis that such new transcription events correspond to biased transcript activity. However, such a correlation between TE overlap and new transcription events suggests that TEs insertion could have contributed to the emergence of the new transcripts that are unrelated to TEs mobility.

When comparing *de novo* transcripts with their non-transcribed homologs, they did not differ in their proportion of TE insertions. Nevertheless, the impact of greater length of TE overlap and higher number of TE insertions seems detrimental for transcription, since *de novo* transcripts have a shorter TE overlap and less numerous TE insertions compared to their homologs. Furthermore, not all TEs seem to impact transcription gain, with RNA transposon being more disruptive than DNA transposon. Indeed, *de novo* transcripts display a higher proportion of DNA transposons compared to their homologs. These results suggest first that *de novo* transcripts emerge in regions that are prone to TE mobility, and are highly variable due to TEs activity. Second, given that DNA TEs are more associated with new transcription events, the insertion of DNA TEs seem to be the more likely to initiate novel transcription. Interestingly, the main difference in TE composition of *de novo* transcripts compared to intergenic sequences, was the higher amount of overlap with retrotransposons (mainly LTR elements from the gypsy family). In *Drosophila melanogaster* certain TEs, such as LTR retrotransposons are reported to be more active than others (Kofler et al., 2015; Petrov et al., 2011). High TE activity can also strongly reshuffle genomes. This could explain why 25% of *de novo* transcripts had no detected transcribed homolog when requiring a high degree (80% identity) of sequence similarity between transcript and homolog. Finally, most of the *de novo* transcripts show high CpGoe values which suggest low DNA methylation, indicating that the genomic location of the transcripts is accessible for the transcription machinery (Roder et al., 2000; Rollins et al., 2006). Furthermore, the correlation between the length of TE overlap with a *de novo* transcript and CpGoe values highlights the impact of TEs bringing along their epigenetic marks.

Taken together, all these independent outcomes strengthen the hypothesis that TEs are actively transposing in *D*.*melanogaster*, and that such activity is noticeable even between lines or individuals. This lines up with previous studies reporting high activity of several TE families in *Drosophila* (Kofler et al., 2015; Bourque et al., 2018; Lawlor et al., 2021; Mérel et al., 2020). Moreover, the significant overlap of active TEs with *de novo* transcripts strongly suggests that TE activity plays a role in initiating new transcription events in intergenic genome regions.

### Minimal tF motifs enrichment leads to transcription gain

Intergenic regions of genomes are known to contain a high proportion of (distal) enhancers which interact with highly distant promoters (Small and Arnosti, 2020). That was confirmed in our results, with random intergenic sequences being the most enriched in highly conserved tF motifs. However, when studying motifs with lower scores of similarity to annotated motifs (80%), *de novo* transcripts contained the highest amount of such motifs, compared to genes and intergenic sequences. Indeed, such low tF motifs, also called sub-optimal transcription factor motifs, appears to be a significant factor for initiating new transcription in genomes. *De novo* transcripts showed lower expression levels than expressed genes, in line with the finding that transcription is initiated at low levels without the presence of strong core motifs (Palazzo and Lee, 2015).

While *de novo* transcripts showed high motifs enrichment of minimum tF motifs, upstream regions of transcripts overlapping with TEs showed the highest amount of low TF motifs. Such enrichment was still lower than in TEs. Most TEs possess a machinery for transcription, which necessitate the presence of tF motifs in their sequence (Chuong et al., 2017). The enrichment of low tF motifs upstream of *de novo* transcripts overlapping with TEs opens two hypothesis. First, the insertion of new TEs in previously untranscribed genomic location could provide sufficient sequence disruption to mutate into minimum tF motifs. tF motifs are usually shorter than 15 nucleotides, and several position allow nucleotide variability without affecting the binding. Therefore, the possibility of a motif emergence caused by mutations due to TE insertions does not seem unlikely. As a second hypothesis, new transcripts could have benefited from the presence of tF motifs in TEs to initiate new transcription events. While these two hypothesis could find support in literature (Chuong et al., 2017; Moschetti et al., 2020), our data seem to give more credit to the second one. Indeed, low tF enrichment was observed in *de novo* transcript compared to their non-transcribed homologs, only when a TE insertion within the sequence was present. Furthermore, while *de novo* transcripts and their homologs shared similar proportion of TE insertions, the TE content of *de novo* transcripts and their homologs diverge. *De novo* transcripts overlap more with DNA TEs, while non-transcribed homologs overlaps more with RNA TEs. Therefore, if TE insertions were to disrupt genomics sequences, both TE families would be expected to generate a similar amount of disruption, and generate a similar amount of motifs. However, both TEs families do not carry the same tF motifs, as their insertion mechanisms diverge. Indeed, our results tend to suggest that DNA TE insertion generates more new transcription events, and that this could be due to the recycling of their tF motifs.

Many different regulatory elements were shown to have been gained through a TE insertion, such as enhancers/enhancer-like elements, promoters, splice sites, cis-regulatory elements, poly-A signals and more (Moschetti et al., 2020). In non-coding regions, transcription can also be initiated through transposable elements (TE) (Kapusta and Feschotte, 2014). TEs have been shown to have the ability to induce a regulatory sequence trough different mechanisms such as domestication (use of TEs for a new function), gene duplication, change of gene expression, ectopic recombination (Kapusta and Feschotte, 2014; Moschetti et al., 2020; Rizzon et al., 2002). About 75% of human and 68% the mouse lncRNA include at the minimum one (partial) retrotransposon insertion (Kapusta et al., 2013). In humans TEs provided up to 23 % of non redundant transcription start sites and about 30% of poly-A sites of lncRNA. (Ganesh and Svoboda, 2016). In *Drosophila*, TE content has been shown to be high in long noncoding RNA (Ganesh and Svoboda, 2016; Fort et al., 2021), compared to protein coding genes, which would support our results.

Indeed, TEs (and especially DNA families), could have played a (partial) role in the gain of transcription of new transcripts, e.g. by inserting the motifs enabling the start of transcription. Our outcomes demonstrate that this gain of transcription through TEs is a frequent event, and can occur independently in different lines from a same species. Determining how exactly the TEs lead to the transcription of these regions and which elements (poly-A, promoter, enhancer etc.) they contributed to insert would need further investigation and more detailed comparisons between the transcript (and up- and downstream) sequences and their homologous regions in the outgroup lines.

60% of *de novo* transcripts emerged without overlapping with TEs. These transcripts showed higher minimal tF enrichment than control sequences, but the difference was less obvious than for transcripts overlapping with TEs. Such small enrichment could be explained by the emergence of low identity tF motifs by other mechanisms than TE insertions, like indels, or other sequence reshuffling that we did not investigate, e.g. genomic inversions or duplications. Furthermore, the high GpC content in all *de novo* transcripts could be associated with low methylation, even though genomes methylation is less observed in invertebrate genomes than vertebrates (Klughammer et al., 2023). Also, we found surprisingly low amounts of core promoter motifs upstream *de novo* transcripts. If such motif enrichment was suspected to be lower than upstream genes, it was surprising to find them less enriched than control intergenic sequences. So far, we have no hypothesis for such an output, but it might also play a role in new transcripts emergence.

## Conclusion

Overall, our study reveals the importance of TEs in transcription gain and loss. At a large scale, a high TE density seems to enable transcription, most likely through changes of chromatin organization (Lawson et al., 2023), as TE density was correlated with *de novo* transcripts density within 100 kb windows. At a finer scale, insertions of TEs seems to lead to different outcomes depending on their insertion patterns. Indeed, a singular insertion of DNA transposon shortly overlapping with the transcript sequence tends to favour the gain of transcription, most likely through enrichment of the upstream region with minimal tF motifs. On the contrary, insertions of RNA transposons likely lead to transcription loss, at the exception of multiple RNA transposon insertions accompanied with an enrichment of minimal core promoter in the upstream region.

## Methods

### Detection of de novo transcripts and their non-transcribed homologs

To investigate the molecular mechanisms enabling new transcript emergence, we searched for *de novo* transcripts and their non-transcribed homologs in the transcriptomes and genomes, respectively, of seven lines of *D. melanogaster*, six inbred european lines and one from Zambia (NCBI Bioproject PRJNA929424)(Grandchamp et al., 2022). Transcripts were defined as being *de novo* (i.e. newly emerged) if they met our four criteria: i) detected in one or several of the seven inbred line transcriptomes with a TPM value (transcripts per million) above 0.5(Grandchamp et al., 2023a); ii) no homology to any other annotated transcripts (cRNA and ncNRA) in the *D. melanogaster* reference transcriptome (Table 1); iii) no homology with annotated transcripts (cRNA and ncRNA) of eleven outgroup *Drosophila* and five Diptera species (Table 1); iv) no overlap of transcript genome location with TEs greater than 80%.

**Table 1:** List of reference species used to build the reference database for the blast search.

|  | Species | Accession number | Assembly |
| --- | --- | --- | --- |
| 1 | *Aedes aegypti* | GCA_002204515.1 | AaegL5 |
| 2 | *Anopheles sinensis* | GCA_000472065.2 | AsinS2 |
| 3 | *Culex quinquefasciatus* | GCA_000209185.1 | CpipJ2 |
| 4 | *Drosophila ananassae* | GCA_000005115.1 | dana_caf1 |
| 5 | *Drosophila erecta* | GCA_000005135.1 | dana_caf1 |
| 6 | *Drosophila grimshawi* | GCA_000005155.1 | dgri_caf1 |
| 7 | *Drosophila melanogaster* | GCA_000001215.4 | BDGP6.32 |
| 8 | *Drosophila mojavensis* | GCA_000005175.1 | dmoj_caf1 |
| 9 | *Drosophila persimilis* | GCA_000005195.1 | dper_caf1 |
| 10 | *Drosophila pseudoobscura* | GCA_000001765.2 | Dpse_3.0 |
| 11 | *Drosophila sechellia* | GCA_000005215.1 | dsec_caf1 |
| 12 | *Drosophila simulans* | GCA_000754195.3 | ASM75419v3 |
| 13 | *Drosophila virilis* | GCA_000005245.1 | dvir_caf1 |
| 14 | *Drosophila williston* | GCA_000005925. | dwil_caf1 |
| 15 | *Drosophila yakuba* | GCA_000005975.1 | dyak_caf1 |
| 16 | *Megaselia scalaris* | GCA_000341915.1 | Msca1 |
| 17 | *Teleopsis dalmanni* | GCA_002237135.2 | ASM223713v2 |

Nucleotide BLAST (version 2.12) (Altschul et al., 1990) with the plus option was used to assess homology between inbred *Drosophila melanogaster* lines and reference transcripts. The lack of homology was defined if a transcript did not return a BLAST hit (with a threshold E-value of 0.05), as well as none of its splicing variant.

Bedtools (version 2.3, intersect with default parameters) (Quinlan and Hall, 2010) was used to map *de novo* transcripts onto their respective genome. *De novo* transcripts overlapping with a gene in sense or antisense direction were filtered out, keeping only intergenic *de novo* transcripts.

To better understand the frequency of transcription gain and loss, we quantified the amounts of *de novo* transcripts shared across inbred *D. melanogaster* lines. To that end, a BLAST search (plus strand option, E-value of 0.05) of our *de novo* transcripts were performed against the transcripts of the other lines. Transcripts were deemed to be homologous if they met those three criteria: i) the transcription start sites of transcripts match up in a 200 nucleotide window; ii) the transcription termination sites of transcripts match up in a 200 nucleotide window; iii) transcripts share at least 80% identity.

To precisely categorize the mechanisms underlying the gain of transcription, direct comparisons of the same nucleotide sequences exhibiting different transcription status is mandatory. We, therefore, used *de novo* transcripts, which were not found across all lines, and their location onto their respective genome to find their ‘non-transcribed homologs’. The unspliced sequences of those *de novo* transcripts were retrieved using bedtools (get fasta with the -s option)(Quinlan and Hall, 2010). Those unspliced sequences were then used to identify similar/identical nucleotide sequences in the genome of other lines, which do no posses this *de novo* transcript, using a nucleotide BLAST search (default settings, E-value cut-off 0.05) (Altschul et al., 1990). A nucleotide sequence was defined as a ‘non-transcribed homologs’, if BLAST hits had 80% query coverage with the *de novo* transcript. If a transcript had multiple ‘non-transcribed homologs’ in the same line, only the nucleotide sequence with the lowest E-value, highest percent identity and highest query coverage, was retained.

Non-transcribed homologs were searched per transcript instead of per orthogroup. The original dataset was reduced to avoid confusion i) Alternative spliceforms were reduced to one spliceform per orthogroup; ii) Orthogroup containing lines duplication were removed (iii) All orthogroup member and non-transcribed homologs have their initiation and termination positions in a same window (+/- 200 nt).

### The role of transposable elements in gain of transcription

To unravel the importance of transposable elements in the emergence of *de novo* transcripts, *de novo* annotations of TEs were performed in each inbred line, using the reasonaTE pipeline from the TransposonUltimate software (Riehl et al., 2022). This pipeline was chosen as it combines, compiles, and filters TE annotations from 13 tools with different annotation approaches (Riehl et al., 2022). *De novo* TE annotations of each *D. melanogaster* line genome was used to infer their relative overlap with *de novo* transcripts, as well as with their upstream and downstream regions, with ‘non-transcribed homologs’ and their upstream regions, and as a control with random intergenic regions of 1100 bp length obtained using bedtools (Quinlan and Hall, 2010). Relative overlap was calculated by dividing the overlap length between a sequence and a TE obtained with bedtools (Quinlan and Hall, 2010) with the full of length of the sequence. Up- and downstream regions were defined as 1000 bp length before and after a given sequence, respectively, with a 100 bp overlap with the given sequence (for a total of 1100 bp length). A given sequence could overlap with more than one TE, in this case relative overlap was calculated using all overlapping TEs, and the number of TEs as well as their class and family were calculated.

Moreover to evaluate features associated with gain of transcription at the genome scale, the distribution of *de novo* transcripts and TEs density within a 10kb sliding window, as well as CpGoe (a proxy for DNA methylation), were plotted along chromosomes for each *D. melanogaster* line, using an R script adapted from (Ylla et al., 2021) https://github.com/guillemylla/Crickets_Genome_Annotation.

## Motif enrichment and gain of transcription

### Motif datasets

The presence of specific DNA motifs before a gene is a major factor enabling transcription, we therefore searched for such motif enrichment upstream of *de novo* transcripts and control sequences, using custom python scripts along the Bio-python motifs (Cock et al., 2009) package. To that end, two motifs databases were downloaded as position frequency matrices (PFM) from JASPAR: the JASPAR Core insects (non redundant) database (Castro-Mondragon et al., 2022) and the JASPAR Pol II database (Fornes et al., 2020), containing 146 tF motifs of *D. melanogaster* and 13 core promoter motifs, respectively. While the JASPAR Core insects database was used to find general promoter and proximal enhancer motifs, the JASPAR Pol II database was restricted to the main core promoter motifs. PFM were used to calculate for each motif a position weight matrix (PWM). The PWM was then used to determine a position specific scoring matrix (PSSM). The absolute score of the PSSM was used to calculate the relative score of motif identity (Formula 1), which was then used as a threshold to determine motifs enrichment. Motifs with a relative score of motif identity superior or equal to 0.8 to the PFM were considered to be enriched in a given sequence. Two types of motifs enrichment were defined: high similarity motif enrichment when a motif had a score above 0.95 and low similarity motif enrichment when its score was between 0.8 and 1. Motif enrichment was estimated for upstream sequences (1000 bp before transcript start and 100 bp after it) of *de novo* transcript, for upstream sequences of ‘non-transcribed’ homologs and for random intergenic sequences of 1100 bp (obtained with bedtools (Quinlan and Hall, 2010), N = 53,300) as negative controls, and for upstream sequences of annotated genes as a positive control. We also restricted our upstream sequences, to 200 bp before a given sequence start and 100 bp after it, to estimate the core promoter binding motifs enrichment as those motifs are expected to be closer to the start of a transcript than general promoter and proximal enhancer (Butler and Kadonaga, 2002).

Formula 1:

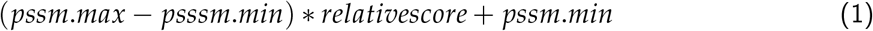

Relative score:

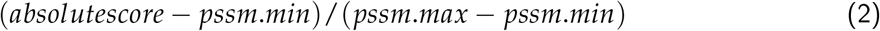

All comparisons of transcripts with other sequence types were performed using Generalized Linear Mixed Models (GLMMs) using the package glmmTMB (Magnusson et al., 2017), retaining best model after simplifying model with a step-wise factor deletion.

### Transcripts vs. non-transcribed homologs

To unravel the differences among sequences leading to transcription, four GLMMs were built using a binomial distribution. The first one assessed the importance of TE overlap, number and presence / absence in gaining transcription. This model includes as a dependent variable the type of sequence (transcript or ‘non-transcribed’ homolog), as fixed factors the relative overlap with TEs, the number of overlapping TEs, the presence or absence of overlapping TEs, the regions of the sequence (upstream, sequence, downstream), and their interactions. Moreover to account for pseudo-replication, the orthogroup ID of the sequence (single ID shared among transcript and non-transcribed homologs) and *D. melanogaster* line were added as random variables into the GLMM. A second model to account only for motifs enrichment was built with as fixed factors the number of minimal and otpimal tF motifs and the minial and optimal number of core promoter. A third model to account simultaneously for TEs and the different motifs was built by adding as fixed factor the number of the different motifs (motifs, cores, low and high). Finally, a fourth model was built to disentangle the impact of different TE classes (DNA vs. RNA transposon) on transcript and non-transcribed homologs, by adding the TE class as a fixed factor.

### Transcripts vs. genes and intergenic regions

To understand how transcripts differ from genic and intergenic regions, three GLMMs were built. The first GLMM compares the relative overlap of transcripts with TEs with the different sequence types, using a zero-inflated Gamma distribution and as dependent variable: the sequence type, as fixed factor: the relative of overlap with TEs, and a random variable: the *D. melanogaster* line. The second GLMM compares the sequence types in term of motif numbers, using a poisson distribution and as dependent variable: the number of motifs / cores, as fixed factor: the sequence type, and a random variable: the *D. melanogaster* line. The third GLMM accounts for differences of sequence features among the different sequence types, using a zero-inflated Gamma distribution and as dependent variable: the sequence type, as fixed factor: the log TPM, the GC content, spliced length, and exon number, and a random variable: the *D. melanogaster* line. As the data-sets were of unequal sample size among the different sequence types and to ensure the robustness of our results, p-values of the best GLMM was bootstrapped using data-sets with equal sample size, using the package boot (Canty and Ripley, 2017).

Furthermore, the density of *de novo* transcript per 100 kb was correlated to its distance to the center of the chromosome and the density of TEs, using GLMMs with as a dependent variable the number of *de novo* trasncript within a 100kb window, as a random variables the chromosome and population, and as an explanatory variable the distance from the center of the chromosome (scaled) and the density of TE per 100kb (scaled), repsectively. Furthermore, the levels of CpGoe of *de novo* transcript was correlated with their relatvie overlap with TEs, using a GLMM with CpGoe value as a dependent variable, length of overlap with a TE as explanatory variable and chromosome and population as a random variable.

### Visualisation

All graphs and statistics were created with R version *>* 4.1 (Team, 2022). The packages dplyr (Wickham et al., 2022), tidyverse (Wickham et al., 2019) and data.table (Dowle and Srinivasan, 2021) were used for data preparation. The plots were mainly done with ggplot2 (Wickham et al., 2016) and its extensions ggpubr (Kassambara and Kassambara, 2020).

## Data access

The files containing processed data is available in the Zenodo archive https://doi.org/10.5281/zenodo.8403184, and is referred in the main text as “Supplemental Deposit”. Supplemental figures, information, analyses and models are found in the Supplementary Information (SI). All programs are stored on GitHub (https://github.com/MarieLebh).

## Author contributions

MKL contribution : Data Curation, Formal Analysis, Investigation, Methodology, Writing and Reviewing original draft BF contribution : Formal Analysis, Investigation, Methodology, Writing and Reviewing original draft JS contribution : Data Curation, Formal Analysis EBB contribution : Funding Acquisition, Reviewing original draft AG contribution : Conceptualization, Funding Acquisition, Project Administration, Supervision, Validation, Writing and Reviewing original draft.

