## Supplementary material for "DNA Transposons favour de *novo* transcript emergence through enrichment of transcription factor binding motifs": SI

#### S1 Distribution of *de novo* transcripts in chromosomes

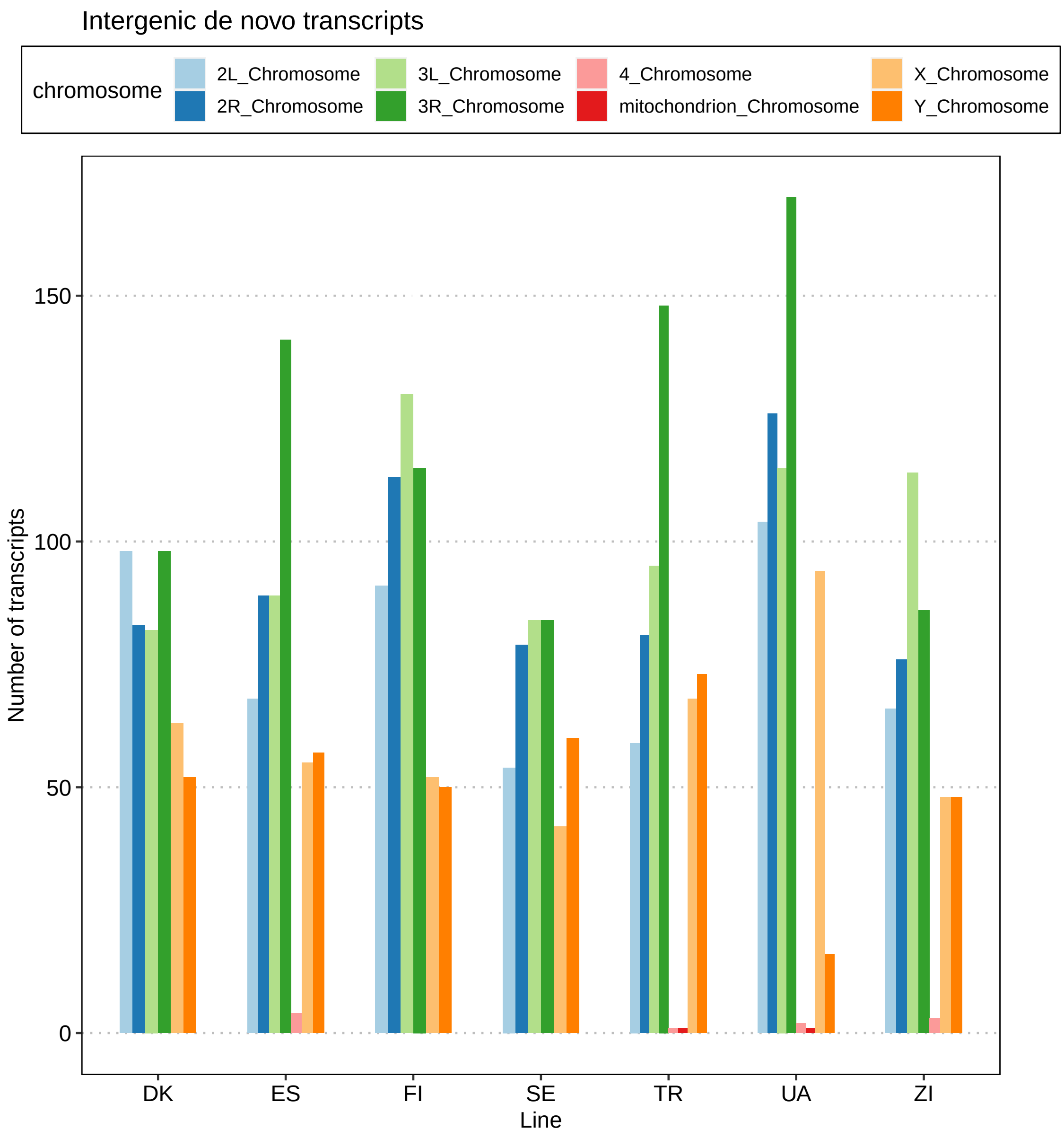

Figure 1: **Number of *de novo* transcripts per chromosomes.** The x axis represents the 7 lines. The y axis represents the number of transcripts emerged *de novo*. Each chromosome arm is represented by a different color.

### S2 Properties of *de novo* transcripts vs annotated transcripts

#### S2.1 Graphs

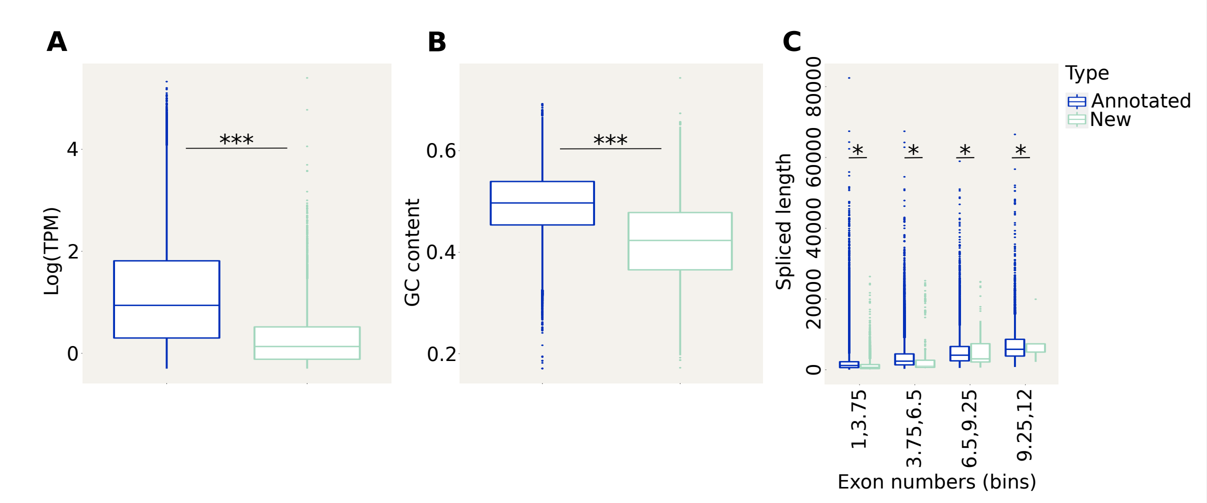

Figure 2: **Properties of *de novo* transcripts and annotated transcripts.** (A): Average TPM values of annotated genic transcripts (dark blue) vs *de novo* transcripts (light blue). (B): Average GC content of annotated genic transcripts vs *de novo* transcripts. (C): Average spliced length of annotated genic transcripts vs *de novo* transcripts, according to the exon/intron number.

#### S2.2 Statistical models

Signif. codes: 0 '***' 0.001 '**' 0.01 '*' 0.05 '.' 0.1 ' ' 1

Test : Anova(modBinAalt,type='III'); Analysis of Deviance Table (Type III Wald chisquare tests)

Response: as.factor(type); Chisq Df Pr(>Chisq)
–> Intercept 286.50 1 < 2.2e-16 ***
–> log10(TPM) 1347.49 1 < 2.2e-16 ***
–> gc_content 2196.85 1 < 2.2e-16 ***
–> spliced_length 405.96 1 < 2.2e-16 ***
–> exon_number 1135.39 1 < 2.2e-16 ***
—

summary(modBinAalt)
Family: binomial ( logit )
Formula: as.factor(type) log10(TPM) + gc_content + spliced_length + exon_number + (1 | chromosome) + (1 | population)

–> AIC BIC logLik deviance df.resid
–> 18139.7 18208.4 -9062.9 18125.7 134943

Random effects:

Conditional model:
Groups Name Variance Std.Dev.
chromosome (Intercept) 0.94561 0.9724
population (Intercept) 0.01494 0.1222
Number of obs: 134950, groups: chromosome, 8; population, 7

Conditional model:
–> Estimate Std. Error z value Pr(>|z|)
–> (Intercept) 6.483e+00 3.830e-01 16.93 <2e-16 ***
–> log10(TPM) -1.350e+00 3.678e-02 -36.71 <2e-16 ***
–> gc_content -1.475e+01 3.146e-01 -46.87 <2e-16 ***
–> spliced_length -1.382e-04 6.857e-06 -20.15 <2e-16 ***
–> exon_number -1.144e+00 3.395e-02 -33.70 <2e-16 ***
—
–> Signif. codes: 0 ‘***’ 0.001 ‘**’ 0.01 ‘*’ 0.05 ‘.’ 0.1 ‘ ’ 1

### S3 Distribution of *de novo* transcripts and TEs along chromosomes

#### S3.1 Visual representation

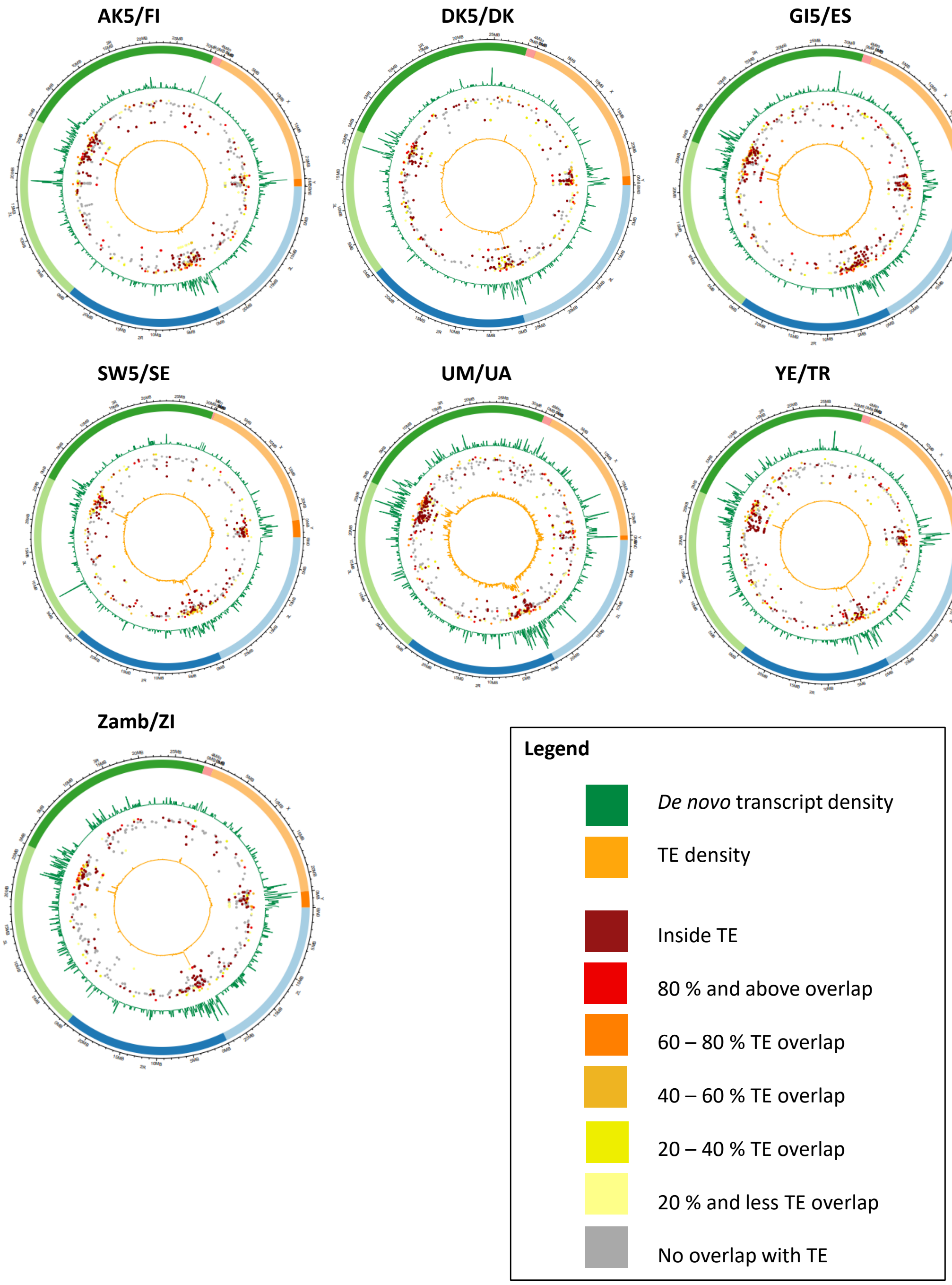

Figure 3

#### S3.2 Figures and statistics of *de novo* transcripts with distance from center

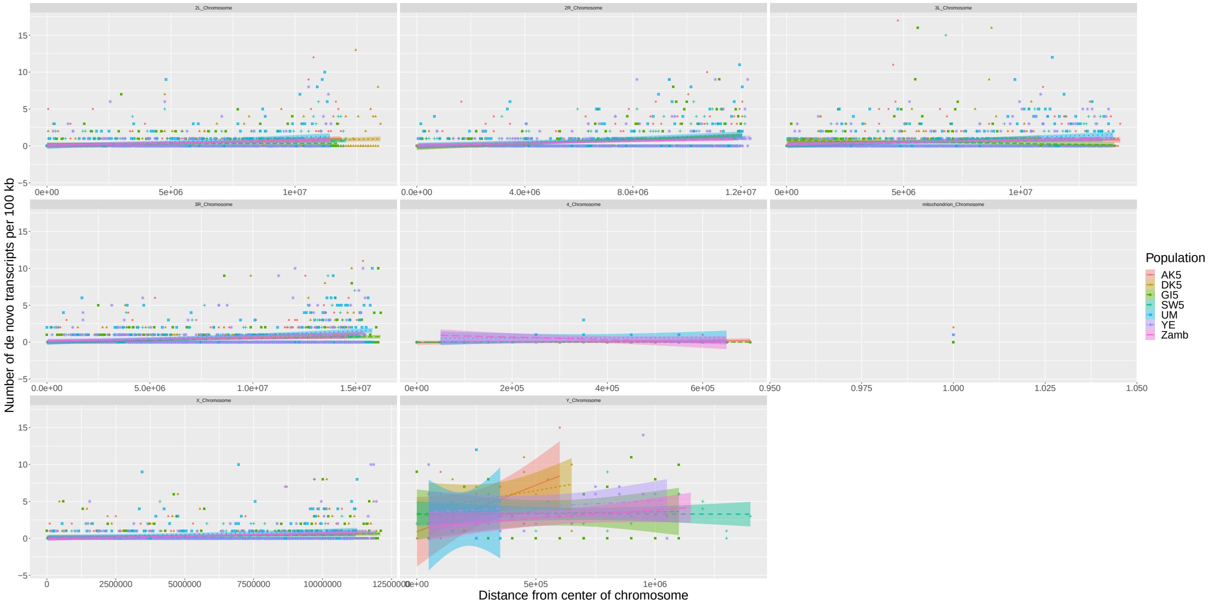

Figure 4: **Linear model *de novo* transcripts distance to the centromere** The x axis represents the distance to the centromere of each chromosome arm. The y axis represents the number of *de novo* transcripts. Each line is represented by a color. The adjusted lines represent the linear models.

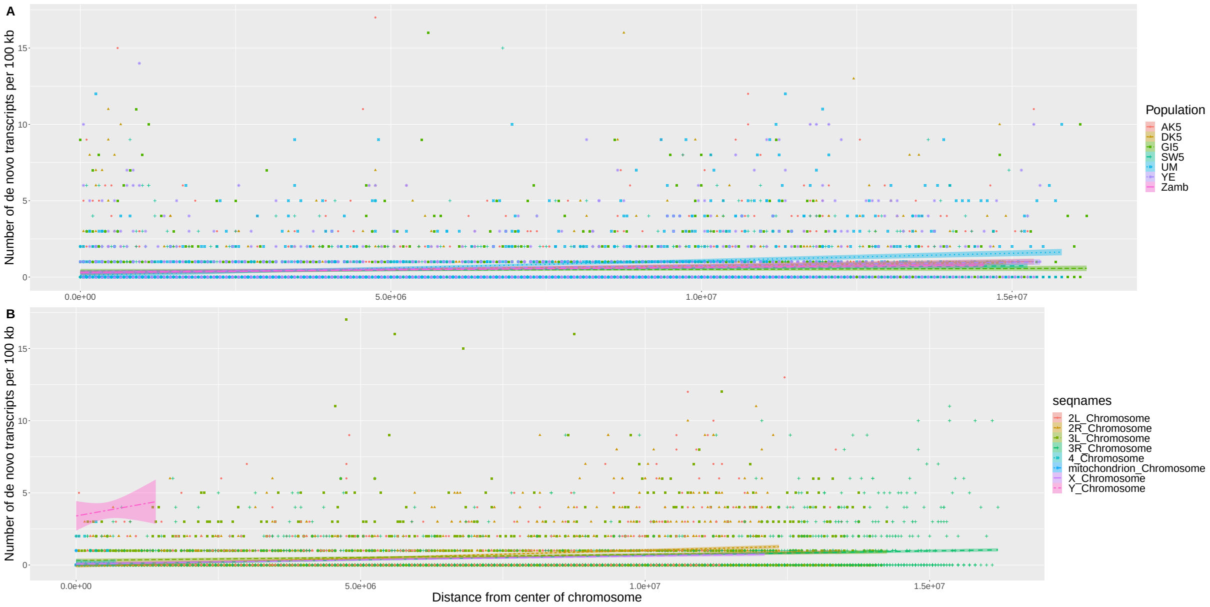

Figure 5: **Linear model *de novo* transcripts distance to the centromere** The x axis represents the distance to the centromere of each chromosome cumulated for (A) and (B). The y axis represents the number of *de novo* transcripts for (A) and (B). (A) Each line is represented by a color. The adjusted lines represent the linear models. (B) Each chromosome is represented by a color.

Signif. codes: 0 ‘***’ 0.001 ‘**’ 0.01 ‘*’ 0.05 ‘.’ 0.1 ‘ ’ 1

–> Anova(mdD,type=’III’)
Analysis of Deviance Table (Type III Wald chisquare tests)

Response: totgenes
–> Chisq Df Pr(>Chisq)
–> (Intercept) 0.9683 1 0.3251
–> distance 973.0866 1 <2e-16 ***
—

> summary(mdD)
Generalized linear mixed model fit by maximum likelihood (Laplace Approximation) [glmerMod]
Family: poisson ( log )
Formula: totgenes   distance + (1 | seqnames) + (1 | Population) Data: df3
Control: glmerControl(optimizer = "bobyqa")

–> AIC BIC logLik deviance df.resid
–> 20559.5 20588.0 -10275.8 20551.5 9233

–> Scaled residuals:
–> Min 1Q Median 3Q Max
–> -2.2909 -0.7382 -0.5667 -0.1996 25.7547

Random effects:
–> Groups Name Variance Std.Dev.
–> seqnames (Intercept) 1.0069 1.0034
–> Population (Intercept) 0.0325 0.1803
–> Number of obs: 9237, groups: seqnames, 8; Population, 7

Fixed effects:
–> Estimate Std. Error z value Pr(>|z|)
–> (Intercept) -0.36133 0.36720 -0.984 0.325
–> distance 0.50205 0.01609 31.194 <2e-16 ***
—

Correlation of Fixed Effects:
(Intr)
distance 0.011

#### S3.3 Figures and statistics of TEs with distance from center

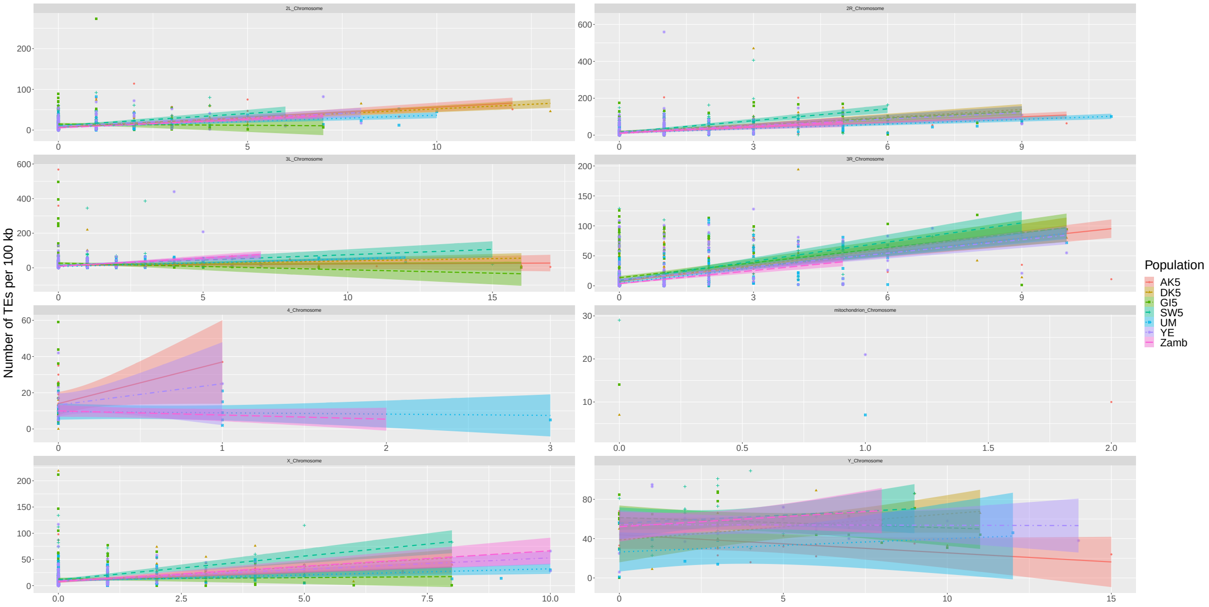

Figure 6: **Linear model TEs distance to the centromere** The x axis represents the distance to the centromere of each chromosome arm. The y axis represents the number of TEs. Each line is represented by a color. The adjusted lines represent the linear models.

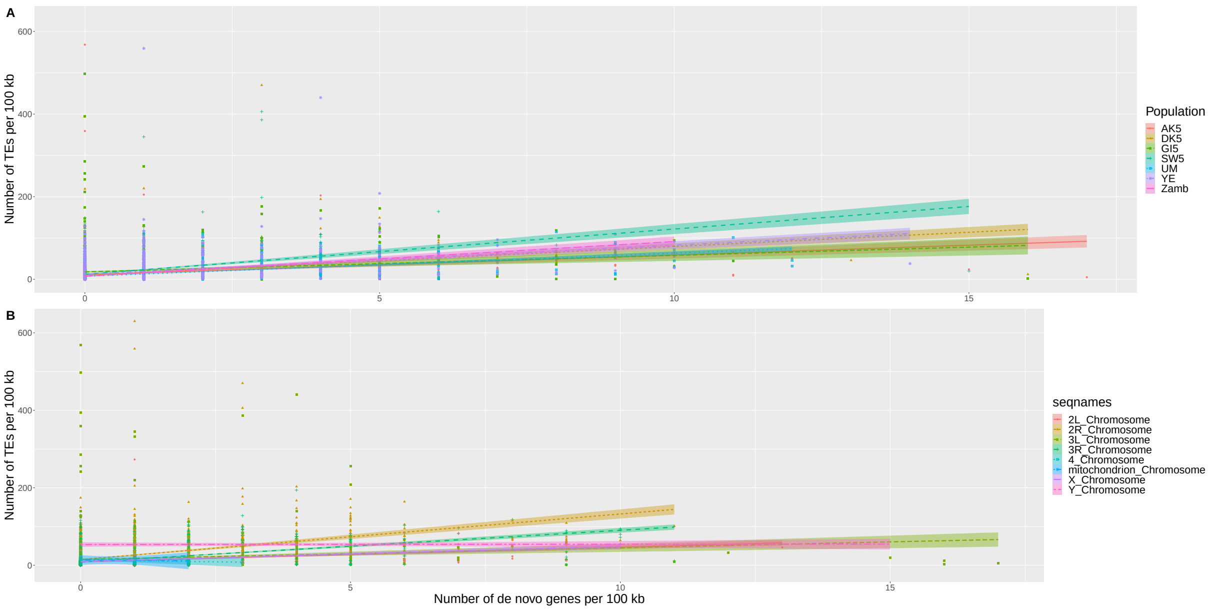

Figure 7: **Linear model TEs distance to the centromere** The x axis represents the distance to the centromere of each chromosome cumulated for (A) and (B). The y axis represents the number of TEs for (A) and (B). (A) Each line is represented by a color. The adjusted lines represent the linear models. (B) Each chromosome is represented by a color.

Anova(mdT,type='III')
Analysis of Deviance Table (Type III Wald chisquare tests)

–> Response: RepCounts
–> Chisq Df Pr(>Chisq)
–> (Intercept) 88.197 1 < 2.2e-16 ***
–> distance 39724.573 1 < 2.2e-16 ***
—
> summary(mdT)
Generalized linear mixed model fit by maximum likelihood (Laplace Approximation) [glmerMod]
Family: poisson ( log )
Formula: RepCounts distance + (1 | seqnames) + (1 | Population) Data: df3
Control: glmerControl(optimizer = "bobyqa")

–> AIC BIC logLik deviance df.resid
–> 197279.3 197307.8 -98635.6 197271.3 9233

–> Scaled residuals:
–> Min 1Q Median 3Q Max
–> -8.522 -2.721 -1.203 1.424 97.218

Random effects:
Groups Name Variance Std.Dev.
seqnames (Intercept) 0.78850 0.8880
Population (Intercept) 0.03327 0.1824
Number of obs: 9237, groups: seqnames, 8; Population, 7

–> Fixed effects:
–> Estimate Std. Error z value Pr(>|z|)
–> (Intercept) 3.008846 0.320386 9.391 <2e-16 ***
–> distance 0.636257 0.003192 199.310 <2e-16 ***

–> Correlation of Fixed Effects:
–> (Intr) distance 0.002

##### S3.4 Statistics Density *de novo* transcripts vs TEs

Anova(md,type='III')
Analysis of Deviance Table (Type III Wald chisquare tests)

–> Response: totgenes
–> Chisq Df Pr(>Chisq)
–> (Intercept) 4.67 1 0.03069 *
–> repcount 1854.77 1 < 2e-16 ***
—
> summary(md)
Generalized linear mixed model fit by maximum likelihood (Laplace Approximation) [glmerMod]

Family: poisson ( log )
Formula: totgenes   repcount + (1 | seqnames) + (1 | Population) Data: df3
Control: glmerControl(optimizer = "bobyqa")

–> AIC BIC logLik deviance df.residu
–> 20702.4 20730.9 -10347.2 20694.4 9233

–> Scaled residuals:
–> Min 1Q Median 3Q Max
–> -5.4304 -0.7115 -0.6387 -0.0429 24.8260

Random effects:
–> Groups Name Variance Std.Dev.
–> seqnames (Intercept) 0.54543 0.7385
–> Population (Intercept) 0.04483 0.2117
–> Number of obs: 9237, groups: seqnames, 8; Population, 7

Fixed effects:
Estimate Std. Error z value Pr(>|z|)
–> (Intercept) -0.604057 0.279524 -2.161 0.0307 *
–> repcount 0.191584 0.004449 43.067 <2e-16 ***
—
Correlation of Fixed Effects:
–> (Intr) repcount -0.010

### S4 Correlation of CpGoe of *de novo* transcripts with TE overlap

#### S4.1 Visual representation

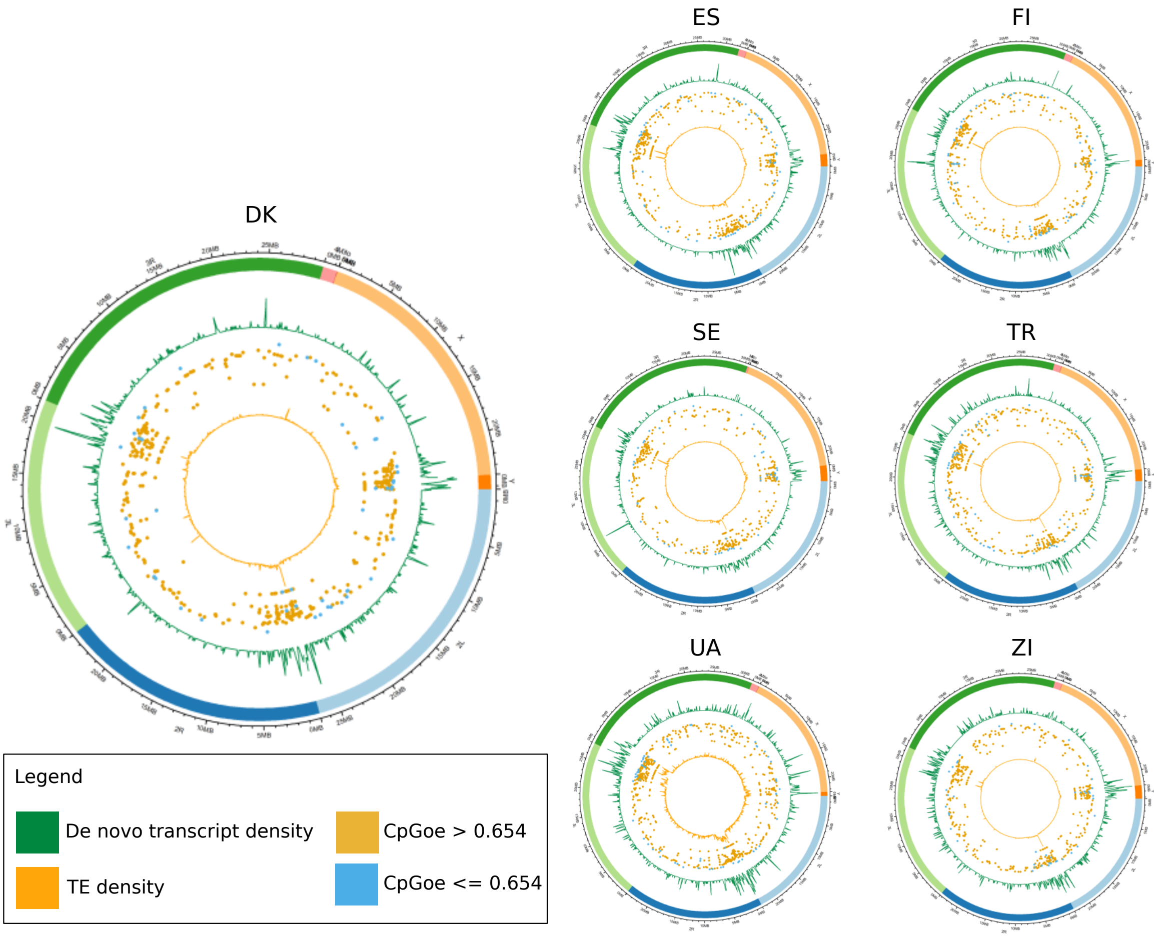

Figure 8:

#### S4.2 Figure correlation

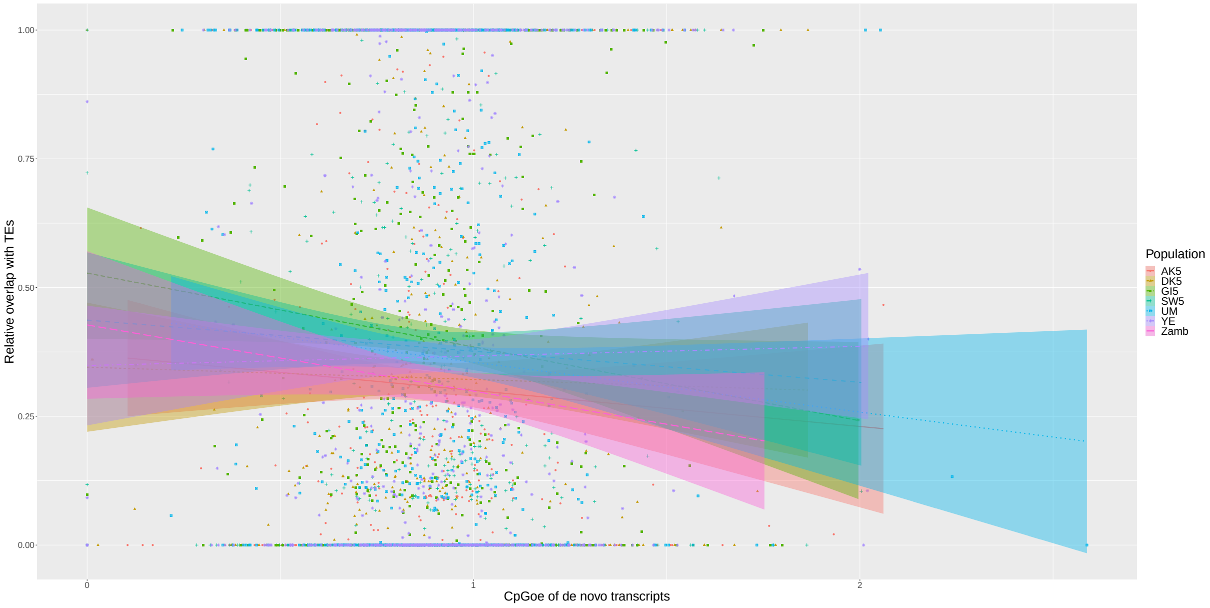

Figure 9: **GC content of *de novo* transcripts according to their TE overlap** The x axis represents the normalised GC content. The y axis represents the average overlap with TEs of *de novo* transcripts. Each lines is represented by a color.

#### S4.3 Statistics of the correlation

Anova(mc,type='III')
Analysis of Deviance Table (Type III Wald chisquare tests)

–> Response: CpGoe
–> Chisq Df Pr(>Chisq)
–> (Intercept) 43408.8216 1 < 2.2e-16 ***
–> TE_ovlp 6.9835 1 0.008226 **
—

> summary(mc)
Linear mixed model fit by REML ['lmerMod']
Formula: CpGoe   TE_ovlp + (1 | Population) + (1 | scaff)
Data: dc
Control: lmerControl(optimizer = "nloptwrap")

REML criterion at convergence: -885.7

–> Scaled residuals:
–> Min 1Q Median 3Q Max
–> -4.1036 -0.5391 -0.0039 0.4760 7.5641

Random effects:
–> Groups Name Variance Std.Dev.
–> scaff (Intercept) 1.952e-05 0.004418
–> Population (Intercept) 0.000e+00 0.000000
–> Residual 4.942e-02 0.222311
–> Number of obs: 5328, groups: scaff, 8; Population, 7

Fixed effects:
–> Estimate Std. Error t value
–> (Intercept) 0.908274 0.004359 208.348
–> TE_ovlp -0.018914 0.007157 -2.643

Correlation of Fixed Effects:
–> (Intr)
–> TE_ovlp -0.576
–> optimizer (nloptwrap) convergence code: 0 (OK)
–> boundary (singular) fit: see help('isSingular')

#### S5 TEs overlapping *de novo* transcripts and non-transcribed homologs

##### S5.1 Figures

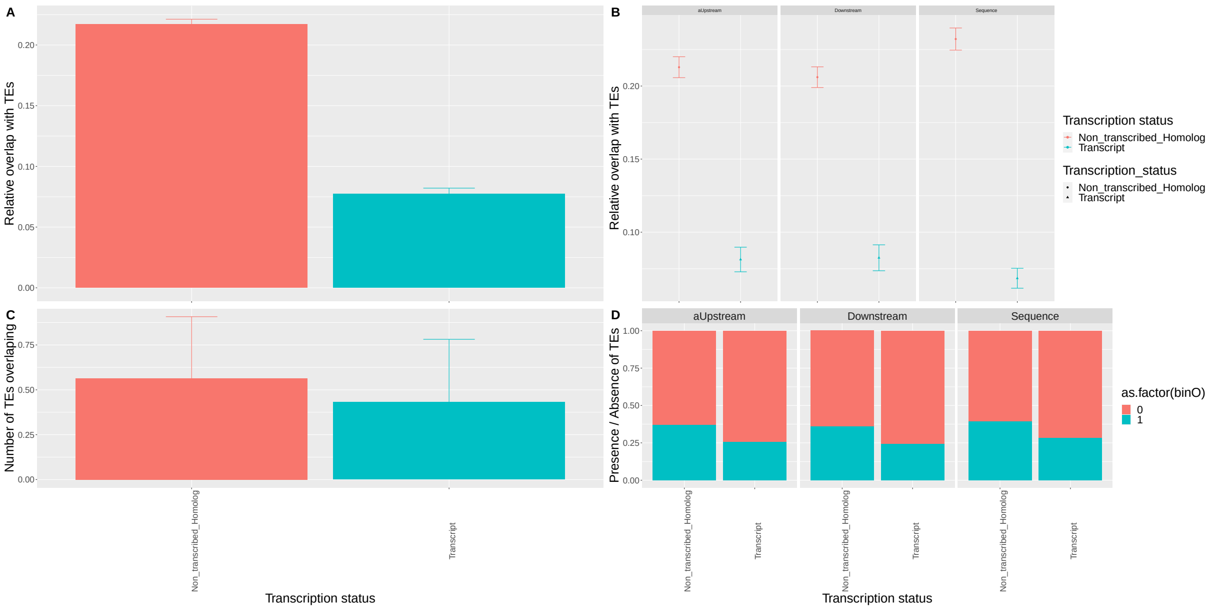

Figure 10: **TEs overlapping *de novo* transcripts and non-transcribed homologs** (A) Relative overlap with TEs. The blue color represents *de novo* transcripts, the pink color represents the non-transcribed homologs. (B) Relative overlap with TEs of upstream, downstream and sequence of *de novo* transcripts and their non-transcribed homologs. The blue color represents *de novo* transcripts, the pink color represents the non-transcribed homologs. (C) Average number of TEs overlapping with : blue : *de novo* transcripts; pink : non-transcribed homologs. (D) Presence/Absence of at least one TE overlap with upstream, downstream and sequences of *de novo* transcripts and their non-transribed homologs.

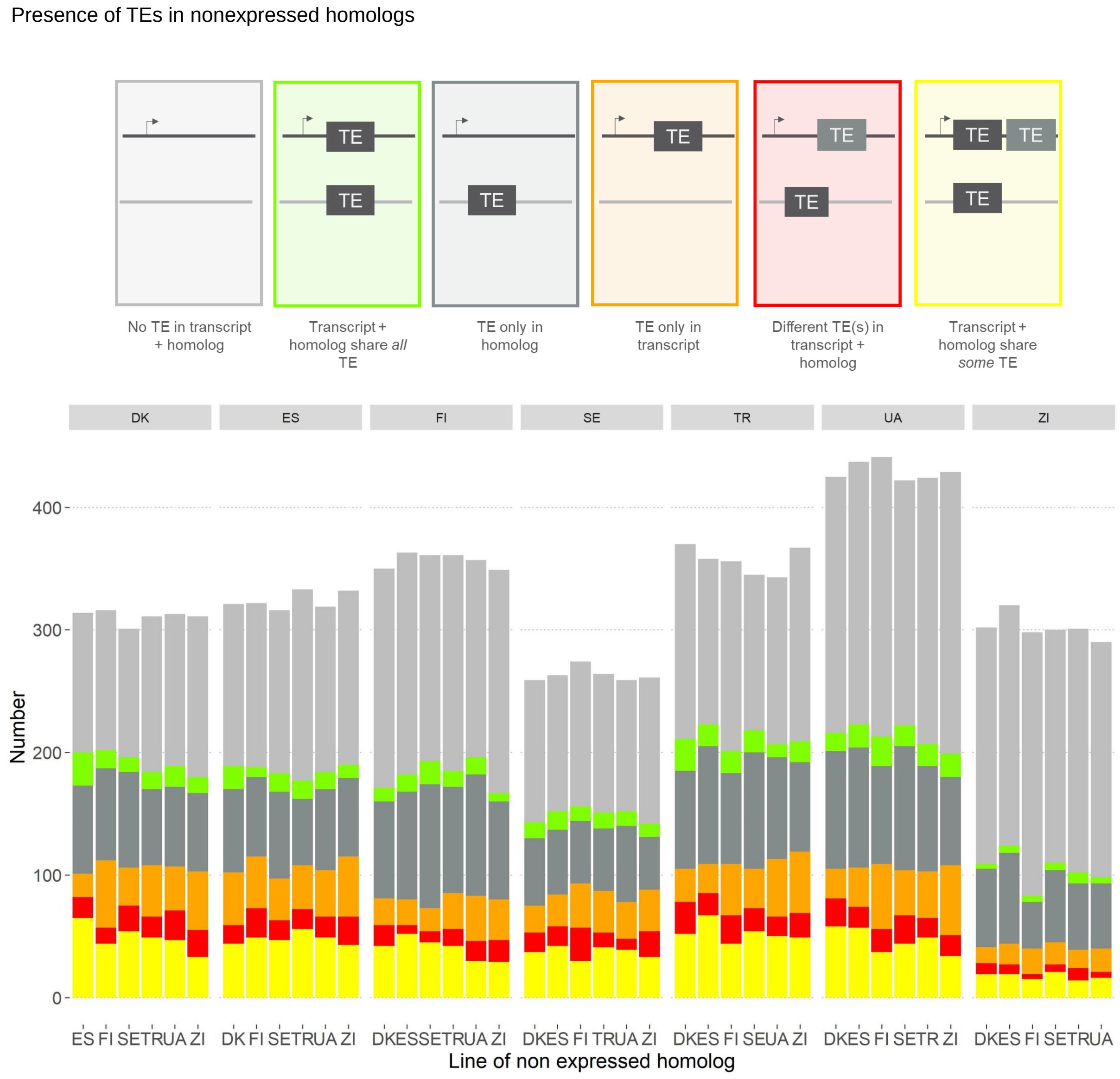

Figure 11: **TEs comparison between *de novo* transcripts and their non-transcribed homologs**. From the left to the right (top to bottom in the plot) : Light grey: Number of *de novo* transcripts and their non-transcribed homologs that both do not overlap with TEs. Green : Number of *de novo* transcripts and their non-transcribed homologs that both overlap the same TE. Grey: Number of *de novo* transcripts that do not overlap with a TE while their non-transcribed homologs do. Orange : Number of *de novo* transcripts that overlap with a TE while their non-transcribed homologs do no. Red : Number of *de novo* transcripts and their non-transcribed homologs that both do overlap with a different TE. Yellow : Number of *de novo* transcripts and their non-transcribed homologs that both do overlap with TEs but some of the overlapping TEs are different.

##### S5.2 Statistics

summary(get.models(msTE1, 1)[[1]])
Family: binomial ( logit )
Formula:
as.factor(Transcription_status) binO + TE_num + TE_ovlp + (1 | Number_Orthogroup) + (1 | Population) + binO:reg + TE_num:reg + TE_ovlp:reg
Data: dataTE

–> AIC BIC logLik deviance df.resid
–> 30462.6 30564.3 -15219.3 30438.6 35547

Random effects:

–> Conditional model:
–> Groups Name Variance Std.Dev.
–> Number_Orthogroup (Intercept) 1.241e-09 3.522e-05
–> Population (Intercept) 4.415e-02 2.101e-01
–> Number of obs: 35559, groups: Number_Orthogroup, 1849; Population, 7

Conditional model:
–> Estimate Std. Error z value Pr(>|z|)
–> (Intercept) -1.478489 0.081240 -18.199 < 2e-16 ***
–> binO 0.071633 0.105015 0.682 0.4952
–> TE_num 0.302268 0.064111 4.715 2.42e-06 ***
–> TE_ovlp -2.345089 0.171987 -13.635 < 2e-16 ***
–> binO:regDownstream -0.007714 0.152170 -0.051 0.9596
–> binO:regSequence 0.284685 0.131719 2.161 0.0307 *
–> TE_num:regDownstream -0.136283 0.093741 -1.454 0.1460
–> TE_num:regSequence -0.160011 0.069320 -2.308 0.0210 *
–> TE_ovlp:regDownstream 0.446712 0.235005 1.901 0.0573 .
–> TE_ovlp:regSequence -0.494534 0.242871 -2.036 0.0417 *
— –> Signif. codes: 0 '***' 0.001 '**' 0.01 '*' 0.05 '.' 0.1 ' ' 1
> Anova(get.models(msTE1, 1)[[1]]) wrong TODO use only type 'III'
Analysis of Deviance Table (Type II Wald chisquare tests)

Response: as.factor(Transcription_status)
Chisq Df Pr(>Chisq)
–> binO 12.1407 1 0.0004933 ***
–> TE_num 51.7351 1 6.352e-13 ***
–> TE_ovlp 583.6019 1 < 2.2e-16 ***
–> binO:reg 6.7192 2 0.0347492 *
–> TE_num:reg 5.3282 2 0.0696608 .
–> TE_ovlp:reg 16.0865 2 0.0003213 ***
—

> Anova(get.models(msTE1, 1)[[1]],type='III')
Analysis of Deviance Table (Type III Wald chisquare tests)

Response: as.factor(Transcription_status)
–> Chisq Df Pr(>Chisq)
–> (Intercept) 331.2009 1 < 2.2e-16 ***
–> binO 0.4653 1 0.4951625
–> TE_num 22.2290 1 2.42e-06 ***
–> TE_ovlp 185.9197 1 < 2.2e-16 ***
–> binO:reg 6.7192 2 0.0347492 *
–> TE_num:reg 5.3282 2 0.0696608 .
–> TE_ovlp:reg 16.0865 2 0.0003213 ***
—

### S6 Low motifs enrichment

#### S6.1 Statistical models

##### *S6.1.1 Core promoter motifs*

Analysis of Deviance Table (Type III Wald chisquare tests)

–> Response: number_core_per_transcript
–> Chisq Df Pr(>Chisq)
–> (Intercept) 1774238 1 < 2.2e-16 ***
–> type_te 476 4 < 2.2e-16 ***
—
–> > summary(modC)
–> Family: poisson ( log )
–> Formula: number_core_per_transcript type_te + (1 | population)
–> Data: dataC

–> AIC BIC logLik deviance df.resid
–> 1193124.2 1193183.8 -596556.1 1193112.2 153461

Random effects:

Conditional model:
–> Groups Name Variance Std.Dev.
–> population (Intercept) 1.926e-11 4.388e-06
–> Number of obs: 153467, groups: population, 7

Conditional model:
–> Estimate Std. Error z value Pr(>|z|)
–> (Intercept) 3.958e+00 2.972e-03 1332.0 <2e-16 ***
–> type_tedn_overlap_TE -1.918e-05 4.781e-03 0.0 0.997
–> type_tedn_TE -2.603e-03 4.412e-03 -0.6 0.555
–> type_tegene 3.772e-02 3.004e-03 12.6 <2e-16 ***
–> type_teIntergenic 2.914e-02 3.030e-03 9.6 <2e-16 ***
—

##### *S6.1.2 Tf motifs*

> Anova(modMI,type='III')
Analysis of Deviance Table (Type III Wald chisquare tests)

Response: number_motifs_per_transcript
–> Chisq Df Pr(>Chisq)
–> (Intercept) 73152613 1 < 2.2e-16 ***
–> type_te 163387 4 < 2.2e-16 ***
— Signif. codes: 0 '***' 0.001 '**' 0.01 '*' 0.05 '.' 0.1 ' ' 1
> summary(modMI)

–> Family: poisson ( log )
–> Formula: number_motifs_per_transcript type_te + (1 | population)
–> Data: dataM

–> AIC BIC logLik deviance df.resid
–> 14947764 14947824 -7473876 14947752 153459

Random effects:

Conditional model:
–> Groups Name Variance Std.Dev.
–> population (Intercept) 1.31e-06 0.001145
–> Number of obs: 153465, groups: population, 7

Conditional model:
–> Estimate Std. Error z value Pr(>|z|)
–> (Intercept) 6.9040836 0.0008072 8553 <2e-16 ***
–> type_tedn_overlap_TE 0.0330909 0.0010855 30 <2e-16 ***
–> type_tedn_TE 0.0602869 0.0009947 61 <2e-16 ***
–> type_tegene -0.0695196 0.0006898 -101 <2e-16 ***
–> type_teIntergenic -0.0066626 0.0006952 -10 <2e-16 ***

#### S6.2 Individual motifs Tf

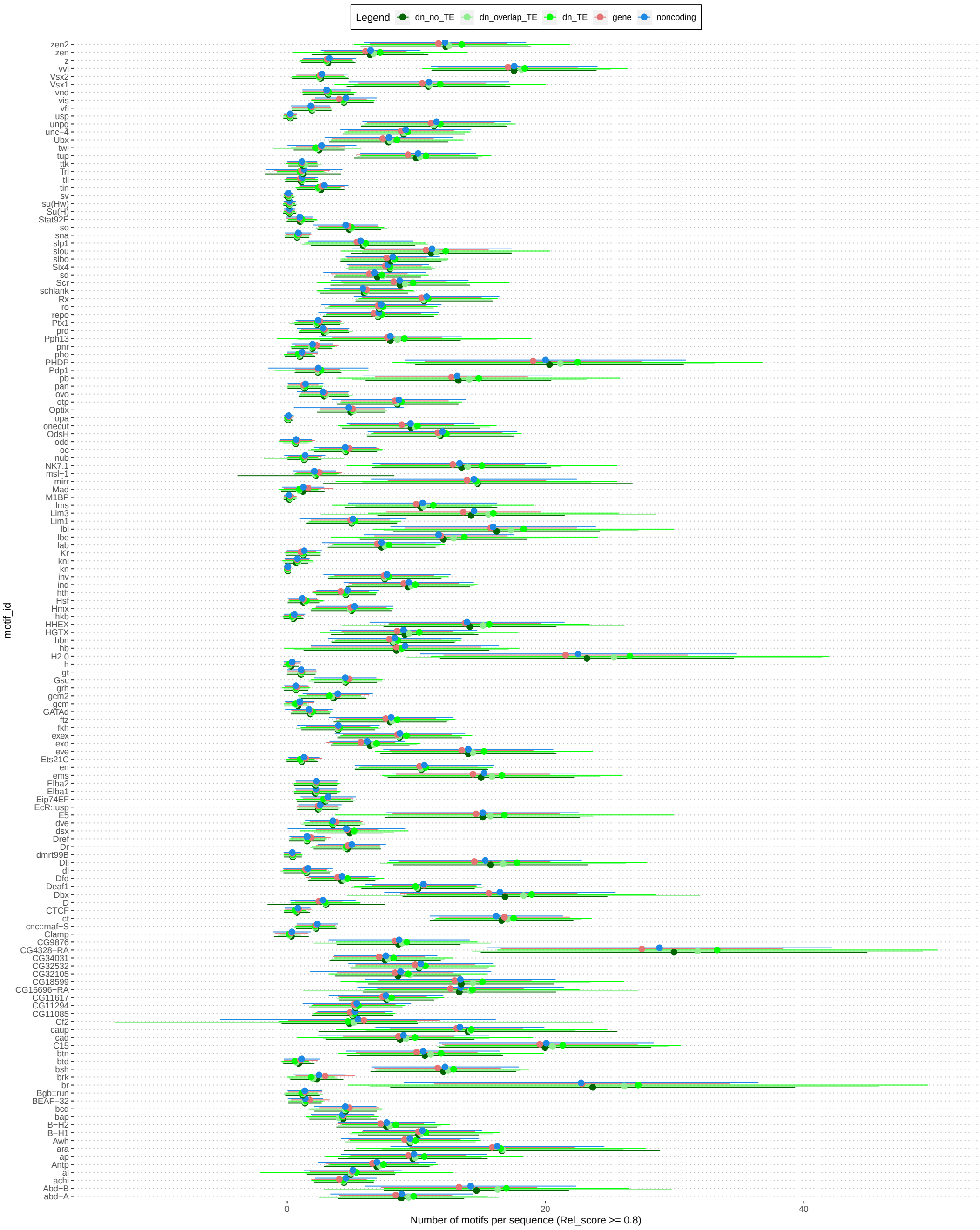

Figure 12: **low tF motifs** Each low tF motif is represented with the average enrichment it show upstrem : *de novo* transcripts without TE overlap, *de novo* transcripts with TE overlap, expressed TEs, genes, random intergenic regions

### S7 High motifs enrichment

#### S7.1 Statistical models

##### *S7.1.1 Core promoter motifs*

Analysis of Deviance Table (Type III Wald chisquare tests)

Response: number_core_high_per_transcript
–> Chisq Df Pr(>Chisq)
–> (Intercept) 7659.1 1 < 2.2e-16 ***
–> type_te 6742.3 4 < 2.2e-16 ***
—
> summary(modCh)
Family: poisson ( log )
Formula: number_core_high_per_transcript type$_te + (1|population)$
$Data : dataC$

–> AIC BIC logLik deviance df.resid
–> 687742.2 687801.8 -343865.1 687730.2 153461

Random effects:

Conditional model:
–> Groups Name Variance Std.Dev.
–> population (Intercept) 2.97e-10 1.723e-05
–> Number of obs: 153467, groups: population, 7

Conditional model:
–> Estimate Std. Error z value Pr(>|z|)
–> (Intercept) 1.090723 0.012463 87.52 < 2e-16 ***
–> type_tedn_overlap_TE -0.005167 0.020082 -0.26 0.79694
–> type_tedn_TE -0.060511 0.018805 -3.22 0.00129 **
–> type_tegene 0.315541 0.012566 25.11 < 2e-16 ***
–> type_teIntergenic 0.099360 0.012690 7.83 4.89e-15 ***
—

##### *S7.1.2 Tf motifs*

Analysis of Deviance Table (Type III Wald chisquare tests)

Response: number_motifs_high_per_transcript
–> Chisq Df Pr(>Chisq)
–> (Intercept) 2053791 1 < 2.2e-16 ***
–> type_te 25735 4 < 2.2e-16 ***
— > summary(modMIh)
Family: poisson ( log )
Formula:

number_motifs_high_per_transcript   type_te + (1 | population)
Data: dataM

–> AIC BIC logLik deviance df.resid
–> 4707456 4707515 -2353722 4707444 153459

Random effects:

Conditional model:
–> Groups Name Variance Std.Dev.
–> population (Intercept) 1.154e-05 0.003396
–> Number of obs: 153465, groups: population, 7

Conditional model:
–> Estimate Std. Error z value Pr(>|z|)
–> (Intercept) 4.199713 0.002930 1433.1 < 2e-16 ***
–> type_tedn_overlap_TE -0.023527 0.004269 -5.5 3.58e-08 ***
–> type_tedn_TE -0.027336 0.003938 -6.9 3.85e-12 ***
–> type_tegene -0.015984 0.002665 -6.0 1.99e-09 ***
–> type_teIntergenic 0.086878 0.002683 32.4 < 2e-16 ***

#### S7.2 Individual motifs Tf

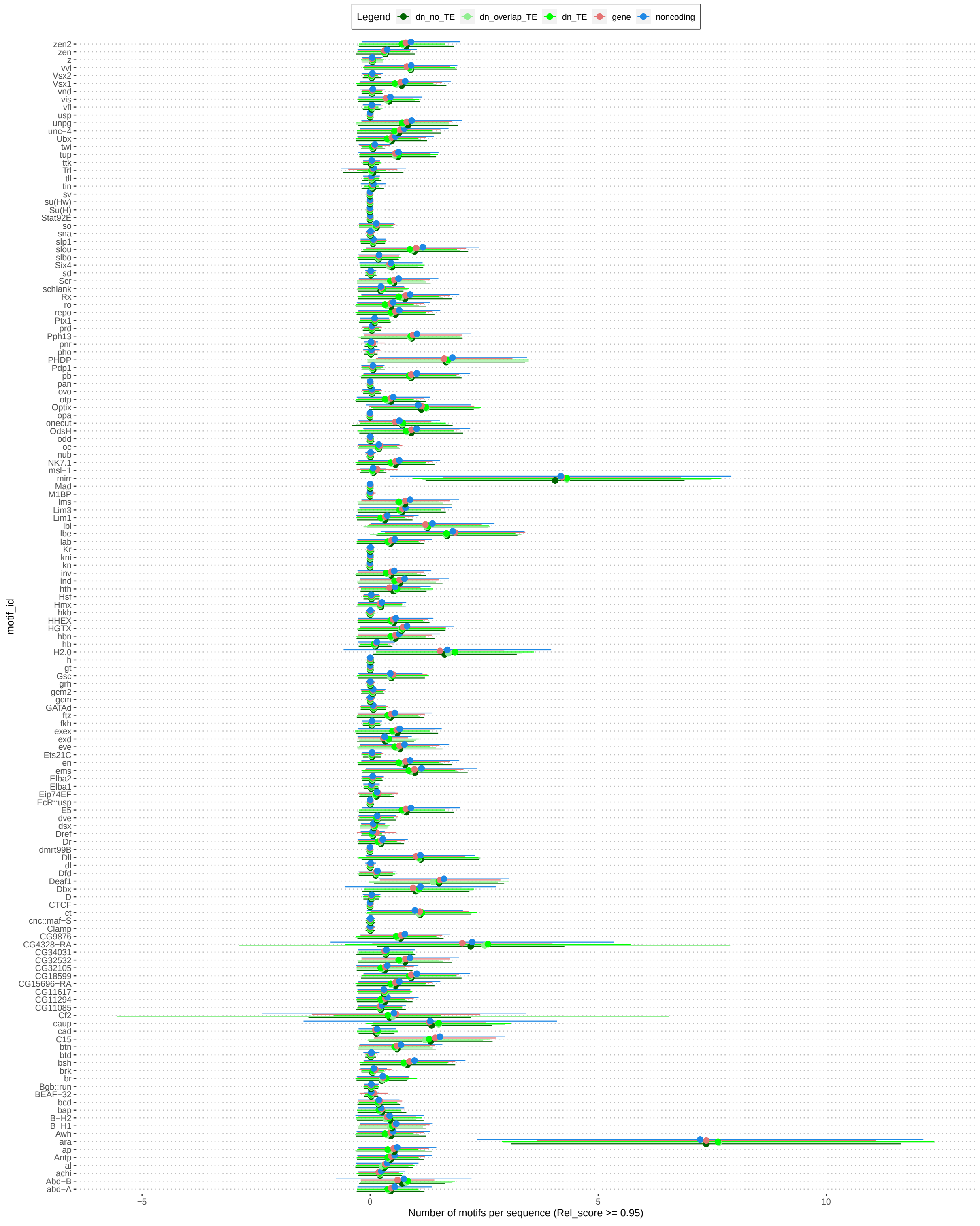

Figure 13: **High tF motifs** Each low tF motif is represented with the average enrichment it show upstrem : *de novo* transcripts without TE overlap, *de novo* transcripts with TE overlap, expressed TEs, genes, random intergenic regions

#### S8 Comparison of motifs enrichment between *de novo* transcripts and non-transcribed homologs

##### *S8.0.1 Global comparison de novo transcripts and non-transcribed homologs*

Anova(modTEMot2,type='III')
Analysis of Deviance Table (Type III Wald chisquare tests)

Response: as.factor(Transcription_status)
–> Chisq Df Pr(>Chisq)
–> (Intercept) 69.7189 1 < 2.2e-16 ***
–> TE_ovlp 278.0631 1 < 2.2e-16 ***
–> TE_num 32.0587 1 1.496e-08 ***
–> Number_core 4.5765 1 0.03241 *
–> binO:Number_motifs 17.7311 1 2.544e-05 ***
—

> summary(modTEMot2)
Family: binomial ( logit )
Formula:
as.factor(Transcription_status)   TE_ovlp + TE_num + Number_core + binO:Number_motifs + (1 | Number_Orthogroup) + (1 | Population)
Data: daTEMot

–> AIC BIC logLik deviance df.resid
–> 9984.7 10036.4 -4985.4 9970.7 11788

Random effects:

Conditional model:)
–> Groups Name Variance Std.Dev.)
–> Number_Orthogroup (Intercept) 1.157e-09 3.401e-05
–> Population (Intercept) 4.286e-02 2.070e-01
–> Number of obs: 11795, groups: Number_Orthogroup, 1849; Population, 7

Conditional model:)
–> Estimate Std. Error z value Pr(>|z|))
–> (Intercept) -1.214e+00 1.453e-01 -8.350 < 2e-16 ***
–> TE_ovlp -2.820e+00 1.691e-01 -16.675 < 2e-16 ***
–> TE_num 1.447e-01 2.556e-02 5.662 1.50e-08 ***
–> Number_core -4.803e-03 2.245e-03 -2.139 0.0324 *
–> binO:Number_motifs 3.236e-04 7.685e-05 4.211 2.54e-05 ***
—

##### *S8.0.2 Comparison de novo transcripts and non-transcribed homologs according to TE class*

Anova(modFCu,type='III')
Analysis of Deviance Table (Type III Wald chisquare tests)

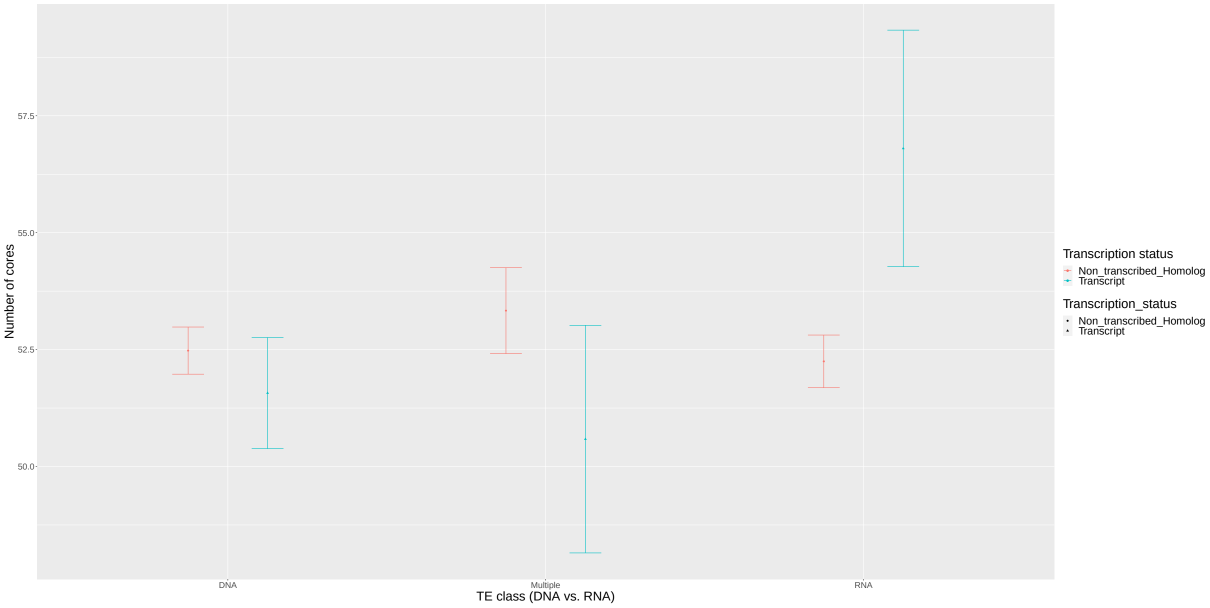

Figure 14: **Comparison *de novo* transcripts and non-transcribed homologs according to TE class.** Blue color represents *de novo* transcripts, pink color represents their non-transcribed homologs. The x axis represents the class of TEs overlapping with the sequences, the y axis represents the number of low core motifs enriched upstream the sequences.

Response: as.factor(Transcription_status)
–> Chisq Df Pr(>Chisq)
–> (Intercept) 14.454 1 0.0001437 ***
–> TE_cla 35.879 2 1.618e-08 ***
–> TE_cla:Number_core 17.566 3 0.0005405 ***
–> TE_cla:TE_num:Number_core 15.423 3 0.0014889 **
— > summary(modFCu)
Family: binomial ( logit )
Formula:
as.factor(Transcription_status) TE_cla + TE_cla:TE_num:Number_core + Number_core:TE_cla + (1 | Number_Orthogroup) + (1 | Population)
Data: OnTE

–> AIC BIC logLik deviance df.resid
–> 3137.6 3208.0 -1557.8 3115.6 4439

Random effects:

Conditional model:
–> Groups Name Variance Std.Dev.
–> Number_Orthogroup (Intercept) 1.184e-09 3.441e-05
–> Population (Intercept) 6.775e-02 2.603e-01
Number of obs: 4450, groups: Number_Orthogroup, 1187; Population, 7

Conditional model:
Estimate Std. Error z value Pr(>|z|)
(Intercept) -1.1560207 0.3040737 -3.802 0.000144 ***
–> TE_claMultiple 0.1471550 0.5721699 0.257 0.797034
–> TE_claRNA -3.6902676 0.6394432 -5.771 7.88e-09 ***
–> TE_claDNA:Number_core -0.0088374 0.0056026 -1.577 0.114705
–> TE_claMultiple:Number_core -0.0297887 0.0101298 -2.941 0.003275 **
–> TE_claRNA:Number_core 0.0291389 0.0115107 2.531 0.011359 *
–> TE_claDNA:Number_core:TE_num 0.0008559 0.0008013 1.068 0.285431
–> TE_claMultiple:Number_core:TE_num 0.0032982 0.0009105 3.623 0.000292 ***
–> TE_claRNA:Number_core:TE_num 0.0045237 0.0041718 1.084 0.278205
